# Oxaliplatin Inhibits RNA Polymerase I via DNA Damage Signaling Targeted to the Nucleolus

**DOI:** 10.1101/2023.04.02.535301

**Authors:** Misha Nechay, Ralph E. Kleiner

## Abstract

Platinum (Pt) compounds are an important class of anti-cancer therapeutics, but outstanding questions remain regarding their mode of action. In particular, emerging evidence indicates that oxaliplatin, a Pt drug used to treat colorectal cancer, kills cells by inducing ribosome biogenesis stress rather than through DNA damage generation, but the underlying mechanism is unknown. Here, we demonstrate that oxaliplatin-induced ribosomal RNA (rRNA) transcriptional silencing and nucleolar stress occur downstream of DNA damage signaling involving ATM and ATR. We show that NBS1 and TOPBP1, two proteins involved in the nucleolar DNA damage response (n-DDR), are recruited to nucleoli upon oxaliplatin treatment. However, we find that rRNA transcriptional inhibition by oxaliplatin does not depend upon NBS1 or TOPBP1, nor does oxaliplatin induce substantial amounts of nucleolar DNA damage, distinguishing it from previously characterized n-DDR pathways. Taken together, our work indicates that oxaliplatin induces a distinct DDR signaling pathway that functions *in trans* to inhibit Pol I transcription in the nucleolus, demonstrating how nucleolar stress can be linked to DNA damage signaling and highlighting an important mechanism of Pt drug cytotoxicity.

## INTRODUCTION

Platinum (Pt) compounds have been a major cornerstone in cancer treatment for nearly 50 years. Cisplatin, the first Pt compound to receive FDA approval in 1978, is the standard of care for the treatment of bladder, lung, and germ cell cancers, among others (Kelland, 2007). To overcome cellular resistance and reduce the toxicity associated with cisplatin, further research has resulted in FDA approval of two additional Pt compounds to date – carboplatin and oxaliplatin, both of which have seen widespread clinical use. Despite the pervasiveness of Pt drugs in cancer treatment, challenges persist in defining their molecular mechanisms of action and predicting clinical outcomes (Rottenberg et al., 2021). For example, colorectal and gastrointestinal cancers were long considered to be insensitive to Pt-based therapeutics (i.e. cisplatin and carboplatin), until oxaliplatin was discovered to be an effective treatment (Raymond et al., 2002). Thus, a better understanding of the relationship between the structure and biological activity of Pt compounds is necessary to improve development and clinical application of this important family of therapeutics.

Most Pt-based drugs in clinical use or investigation contain an electrophilic Pt(II) core which exhibits broad reactivity towards all classes of cellular macromolecules (e.g. DNA, RNA, proteins) (Cunningham and DeRose, 2017; Hostetter et al., 2012). However, the prevailing hypothesis for Pt-induced cytotoxicity only accounts for the accumulation of bulky Pt adducts on DNA, which leads to activation of the DNA damage response (DDR) and pro-apoptotic pathways (Wang and Lippard, 2005). This model, based primarily on studies using the first-generation drug cisplatin, is supported by a wealth of cellular and biochemical data as well as clinical evidence demonstrating that cancers deficient in certain DNA repair factors are particularly sensitive to Pt-based treatments (Rabik and Dolan, 2007). While DNA lesions appear to be the primary mediator of cisplatin cytotoxicity, emerging work has illuminated diverse cellular mechanisms exhibited by Pt compounds containing different substituents around the Pt(II) core (Park et al., 2012; Yonezawa et al., 2006). In particular, phenotypic studies of oxaliplatin have indicated that this compound is better characterized as a transcription/translation inhibitor rather than a canonical DNA damaging agent (Bruno et al., 2017), which has led to the investigation of alternative cellular targets to explain its mechanism of action. Oxaliplatin has further been implicated in the disruption of nucleolar morphology and ribosome assembly (Burger et al., 2010; Sutton et al., 2019), but the underlying mechanism is poorly characterized, and the extent to which different Pt compounds can evoke this response is not well understood.

A number of cellular insults can lead to ‘nucleolar stress’, which is characterized by changes in nucleolar organization, impairment of ribosome biogenesis, and activation of pro-apoptotic signaling through p53 (Yang et al.). One major pathway that induces nucleolar disruption is the nucleolar DNA damage response (n-DDR), a subset of DNA damage signaling in response to ribosomal DNA (rDNA) damage within the nucleolus (Ciccia et al., 2014; Kruhlak et al., 2007). Selective induction of double-strand breaks (DSBs) in rDNA has been shown to inhibit ribosomal RNA (rRNA) synthesis and lead to nucleolar segregation in a manner dependent on the activity of DDR factors such as ATM (Ataxia telangiectasia mutated) and ATR (Ataxia telangiectasia and Rad3-related) kinases, which transduce the DNA damage signal through nucleolar adaptor proteins, such as the ATM-associated phosphoprotein Treacle. Interestingly, rRNA silencing and nucleolar reorganization in response to DSBs occurring outside of nucleoli has also been reported (Larsen et al., 2014), but the mechanism of *in-trans* damage signaling is not well characterized, and the propensity of different DNA damaging agents to induce this effect is unknown. In light of the connection between DNA damage signaling and Pt drugs, and the induction of nucleolar stress by oxaliplatin, an examination of the role of the DDR in oxaliplatin-induced nucleolar disruption and cytotoxicity is warranted.

Here we characterize the mechanism of action of oxaliplatin in human cancer cell lines. We find that in contrast to cisplatin, oxaliplatin is a potent and specific inhibitor of RNA Pol I transcription and rapidly induces nucleolar stress at clinically relevant concentrations. Additionally, we demonstrate that oxaliplatin-induced rRNA silencing and nucleolar disruption occur downstream of ATM and ATR signaling, but in the absence of appreciable nucleolar DNA damage foci. While we observe the recruitment of established n-DDR factors NBS1 and TOPBP1, we show that these do not serve an essential role in transcriptional silencing by oxaliplatin. Taken together, these findings demonstrate how nucleolar stress can be linked to DNA damage signaling and illuminate an important mechanism of oxaliplatin-mediated cytotoxicity.

## RESULTS

### Oxaliplatin is a selective inhibitor of nucleolar transcription

We first set out to characterize the effect of oxaliplatin treatment on nucleolar structure and rRNA transcription. Redistribution of the nucleolar protein NPM1 from the interior of nucleoli to the nucleoplasm is a common indicator of nucleolar stress (Boulon et al., 2010). Using immunofluorescence for endogenous NPM1, we observed predominantly nucleoplasmic distribution of NPM1 accompanied by nucleolar rounding in U2OS cells after 6 hr treatment with 5 µM oxaliplatin (Fig. 1A). These morphological changes were also recapitulated in cells transiently transfected with mCherry-tagged NPM1 (SI Appendix, Fig. S1A). Additionally, we observed the formation of peripheral cap-like structures containing Treacle, a regulator of rRNA transcription (Fig. 1A) (Valdez et al., 2004). These perturbations to nucleoli are consistent with induction of nucleolar stress by oxaliplatin, as reported by Sutton *et al*. (Sutton and DeRose, 2021; Sutton et al., 2019).

**Figure 1.**
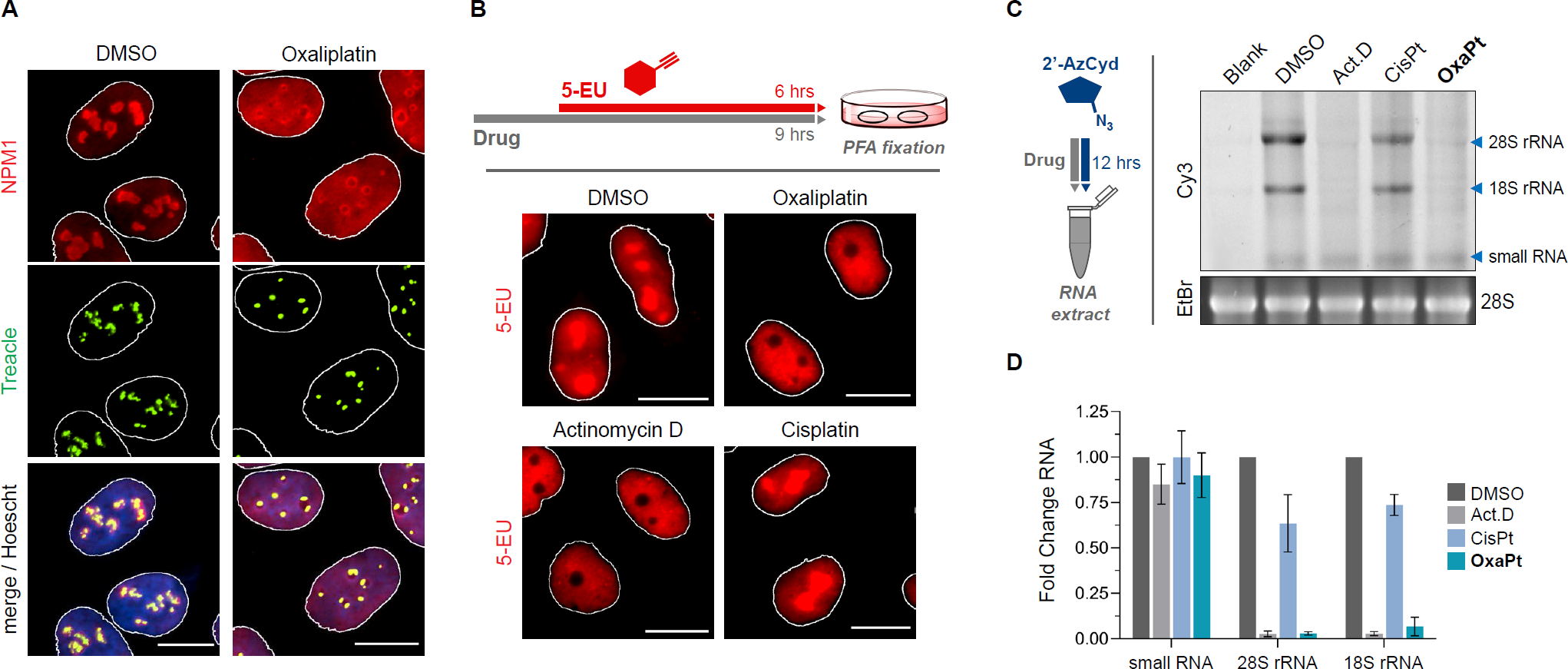
Oxaliplatin selectively inhibits rRNA synthesis and induces nucleolar stress. (A) Representative images of immunostaining against NPM1 (red) and Treacle (green) in U2OS cells with or without (±) treatment of oxaliplatin (5 µM) for 6 hours. (B) Schematic of “long-pulse” 5-EU labeling of bulk nuclear RNA synthesis, and representative images of 5-EU incorporation in fixed HeLa cells after treatment with Pt drugs (20 µM) or Act.D (5 nM) for 9 hours. In (A) and (B), white outlines of nuclei were generated from the DAPI channel. All scalebars = 15 µM. (C) Schematic of 2’AzCyd metabolic labeling of cellular RNA in U2OS cells co-treated with various drugs for 12 hours *(left)*, and detection of extracted RNA on a 1% agarose gel by fluorescence imaging of Cy3-labeled RNA *(right top)* and ethidium bromide staining *(right bottom)*. (D) Quantification of RNA bands in Cy3 channel from (C), relative to signal in DMSO control. Error bars represent mean ± SEM of *N* = 3 biological replicates.

Previous work has shown that rRNA synthesis and processing are closely linked to nucleolar stress and NPM1 redistribution (Boulon et al., 2010). To investigate oxaliplatin’s impact on transcription, we used 5-ethynyluridine (5-EU) to label newly transcribed RNA during drug treatment. Compared to DMSO-treated cells, which showed strong nucleolar 5-EU signal consistent with highly active rRNA transcription, oxaliplatin-treated cells exhibited depletion of fluorescent signal in nucleoli relative to the nucleoplasm (Fig. 1B). A similar pattern was observed with treatment using Actinomycin D (Act.D), a selective inhibitor for Pol I, suggesting that oxaliplatin selectively inhibits rRNA synthesis rather than global RNA transcription. In contrast, 5-EU signal in cisplatin-treated cells was indistinguishable from the DMSO control, indicating that cisplatin has a minimal effect on rRNA transcription. To confirm this finding, we used an alternative approach to metabolically label cellular RNA with 2’-azidocytidine (2’-AzCyd) (Wang et al., 2020), which allowed us to clearly visualize newly transcribed 18S and 28S rRNA bands using in-gel fluorescent detection (Fig. 1C). Co-treatment of U2OS cells with 2’-AzCyd and either oxaliplatin or Act.D for 12 hours resulted in nearly complete abolition of signal from nascent 18S and 28S rRNA, while a faint small RNA band was unchanged in both conditions relative to DMSO-treated cells (Fig. 1C and 1D). Cisplatin, on the other hand, induced a minimal decrease in rRNA transcription under these conditions, consistent with 5-EU labeling results. Together, these data show that oxaliplatin strongly inhibits rRNA synthesis and induces nucleolar stress, distinct from cisplatin.

To investigate the relationship between nucleolar disruption and rRNA synthesis, we simultaneously monitored rRNA synthesis and NPM1 localization in cells treated with different concentrations of oxaliplatin using 5-EU labeling and indirect immunofluorescence (Fig. 2A). We used coefficient of variation (CV) to quantify NPM1 localization, with lower CV values representing a more homogeneous distribution of NPM1 throughout the nucleus indicative of nucleolar stress (Fig. 2B). At 5 µM oxaliplatin treatment, nucleolar transcription and CV for NPM1 are decreased by ∼80% and ∼50%, respectively, compared to untreated cells. Surprisingly, nucleoplasmic redistribution of NPM1 and inhibition of nucleolar RNA transcription could be detected with oxaliplatin treatment concentrations nearly 5-fold lower than the 72 hr IC_50_ in U2OS cells (SI Appendix, Fig. S1B, S1C). By comparison, cisplatin only induces nucleolar stress at dosages 10-fold higher than its IC_50_, a level not considered to be clinically relevant (Burger et al., 2010). Next, we sought to characterize the temporal relationship between nucleolar stress and transcriptional inhibition induced by oxaliplatin. We performed a quantitative time-course analysis of nucleolar transcription and protein localization during a 5 hr oxaliplatin treatment, utilizing a short pulse of 5-EU to detect rRNA synthesis with higher temporal resolution. In addition to analyzing NPM1 distribution, we also measured nucleolar cap formation by counting Treacle foci per cell, which decrease in number as Treacle aggregates on the nucleolar periphery. We observed that NPM1 dispersion and Treacle cap formation begin concurrently in a subset of cells as early as 2 hr of treatment, and both reach steady state within 4 hr (Fig. 2C). We compared this to the kinetics of nucleolar transcription as measured by 5-EU incorporation, which also showed substantial inhibition (∼50% reduction) at 2 hr (Fig. 2D). The similarities in kinetics suggest that Pol I inhibition by oxaliplatin is concurrent with or even slightly precedes nucleolar reorganization.

**Figure 2.**
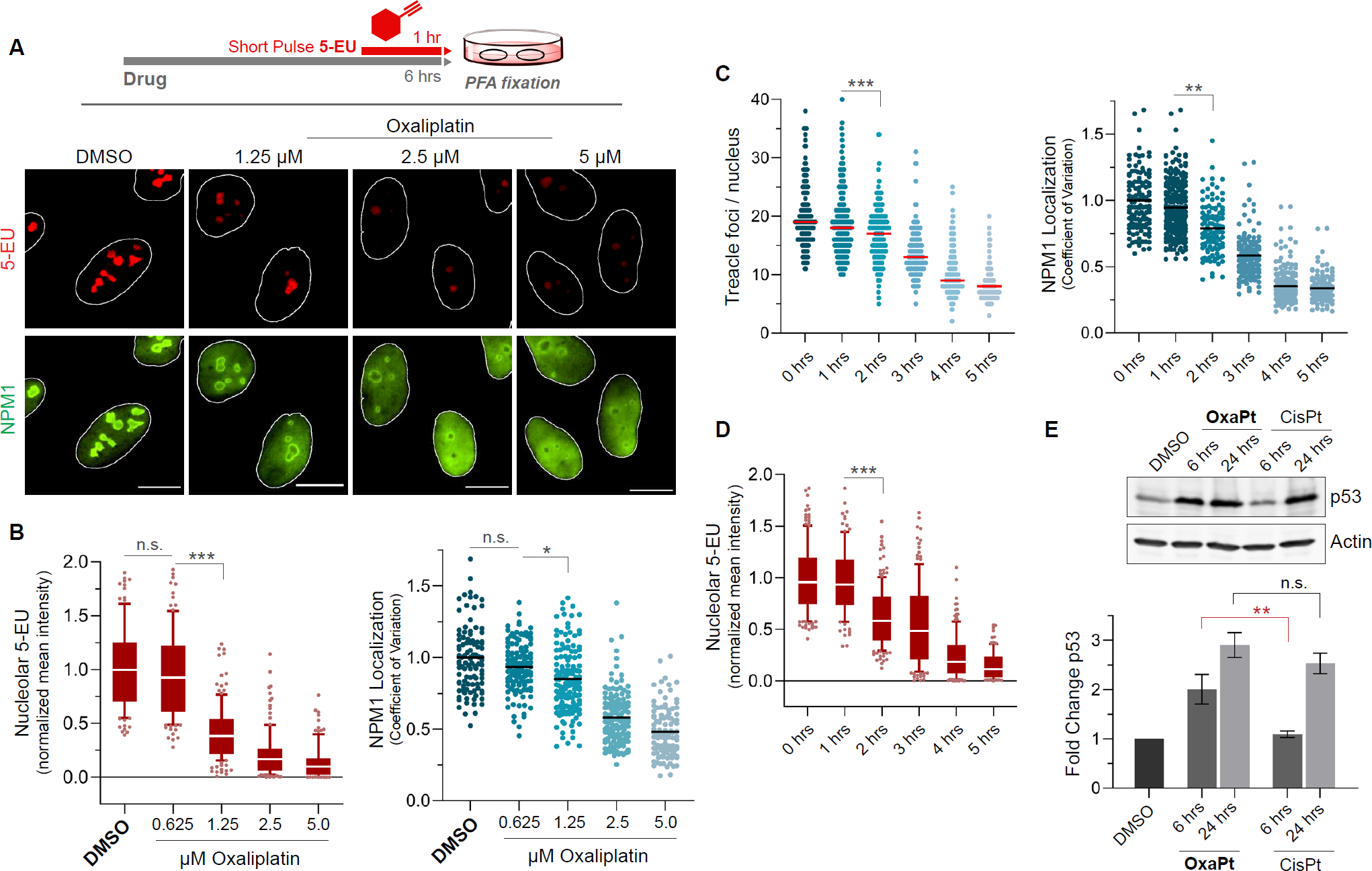
Oxaliplatin is a potent and rapid inhibitor of nucleolar function. (A) Schematic of “short-pulse” 5-EU labeling of nucleolar RNA synthesis *(top)*, and representative images of NPM1 localization and rRNA silencing upon treatment with various concentrations of oxaliplatin for 6 hours *(bottom)*. Scalebars = 15 µM. (B) Quantification of 5-EU signal in nuclei from (A), normalized relative to DMSO control *(left)*. Quantification of NPM1 localization from (A) as measured by the coefficient of variation in NPM1 signal in nuclei *(right)*. Number of cells analyzed for each condition, summed over 2 biological replicates: DMSO (*N* = 103), oxaliplatin at 0.625 µM (*N* = 121), 1.25 µM (*N* = 145), 2.5 µM (*N* = 136), and 5.0 µM (*N* = 104). (C) Quantification of Treacle (TCOF1) foci count per nucleus and NPM1 localization by coefficient of variation in response to treatment with oxaliplatin (5 µM) over a period of 0-5 hours in fixed cells. Number of cells analyzed for each condition, summed over 3 biological replicates: 0 hours (*N* = 129), 1 hour (*N* = 257), 2 hours (*N* = 111), 3 hours (*N* = 141), 4 hours (*N* = 269), 5 hours (*N* = 119). (D) Quantification of rRNA synthesis by nucleolar 5-EU incorporation in cells treated with oxaliplatin (5 µM) over a period of 0-5 hours in fixed cells. Number of cells analyzed for each condition, summed over 2 biological replicates: 0 hours (*N* = 153), 1 hour (*N* = 102), 2 hours (*N* = 172), 3 hours (*N* = 136), 4 hours (*N* = 137), 5 hours (*N* = 96). (E) Western blot of p53 stabilization in response to 10 µM oxaliplatin and cisplatin treatments, with actin loading control underneath *(top)*. Quantification of p53 signal from western blot *(bottom)*. Error bars represent mean ± SEM. White outlines of nuclei in microscopy images were generated from the DAPI channel and used as ROIs for quantifications in (B-D). Statistical analysis in (B-E) was performed using a one-way analysis of variance, Tukey’s multiple comparisons test. Adjusted *p*-values are indicated by ***P<0.001, **P<0.01, *P<0.05, and *n.s.* denotes a non-significant *p*-value > 0.05. All experiments were performed in wild-type U2OS cells.

The inhibition of nucleolar structure and function at early time points led us to question whether this stress response is linked to cell death pathways. The tumor suppressor protein p53 has been shown to mediate pro-apoptotic signaling in response to nucleolar stress, but it is also activated by a multitude of other stressors including DNA damage (Boulon et al., 2010; Yang et al., 2016). Previous work has linked p53 stabilization to DDR induced by cisplatin, and no significant difference was found in p53 levels between prolonged treatment with either cisplatin or oxaliplatin, possibly due to the nonspecific activation of several different stress pathways (Sutton and DeRose, 2021). We measured p53 levels by western blotting and indeed saw similar increases in p53 at 24 hr treatment with either cisplatin or oxaliplatin (Fig. 2E). However, when we examined p53 levels at 6 hr, shortly after the redistribution of NPM1 and rRNA silencing as described earlier, we observed a 2-fold higher level of p53 in oxaliplatin-treated cells compared to those treated with cisplatin. To our knowledge, this evidence of early apoptotic signaling following nucleolar stress induction has not been previously characterized for oxaliplatin, and it suggests a direct connection between nucleolar stress and oxaliplatin cytotoxicity.

### Oxaliplatin binds nucleolar nucleic acids and proteins with lower efficiency than cisplatin

Previous studies have demonstrated that damage to ribosomal DNA can lead to Pol I inhibition and redistribution of nucleolar proteins (Korsholm et al., 2020; Larsen et al., 2014; Lee et al., 2005). Given the propensity of Pt ions to react with guanine-rich nucleic acids (Fichtinger-Schepman et al., 1985) and the precedent for cisplatin to accumulate on ribosomal RNA in yeast (Hostetter et al., 2012), we reasoned that oxaliplatin may form bulky adducts on rRNA or rDNA that are capable of disrupting interactions with transcription factors and other nucleolar proteins. To investigate this, we first analyzed Pt accumulation on rRNA using reverse transcription (RT) primer extension on RNA isolated from U2OS cells treated with either oxaliplatin or cisplatin (Rijal and Chow, 2007). We chose to probe a solvent-accessible region on the 28S rRNA adjacent to the peptidyl transferase center (PTC) and containing known reactive sites for cisplatin and ribosomal inhibitors (Myasnikov et al., 2016; Plakos and DeRose, 2017). We were unable to clearly visualize oxaliplatin adducts in this region (Fig. 3A), as indicated by lack of additional RT stops compared to DMSO-treated sample. In contrast, several dose-dependent cisplatin adducts appeared in the vicinity of the E-site, which may explain cisplatin’s inhibitory effects on protein synthesis (Rosenberg and Sato, 1993). Primer extension probing of 18S and 5S rRNA showed a similar trend with relatively little accumulation of oxaliplatin compared with cisplatin (SI Appendix, Fig. S2), suggesting that rRNA is not a major target of oxaliplatin.

**Figure 3.**
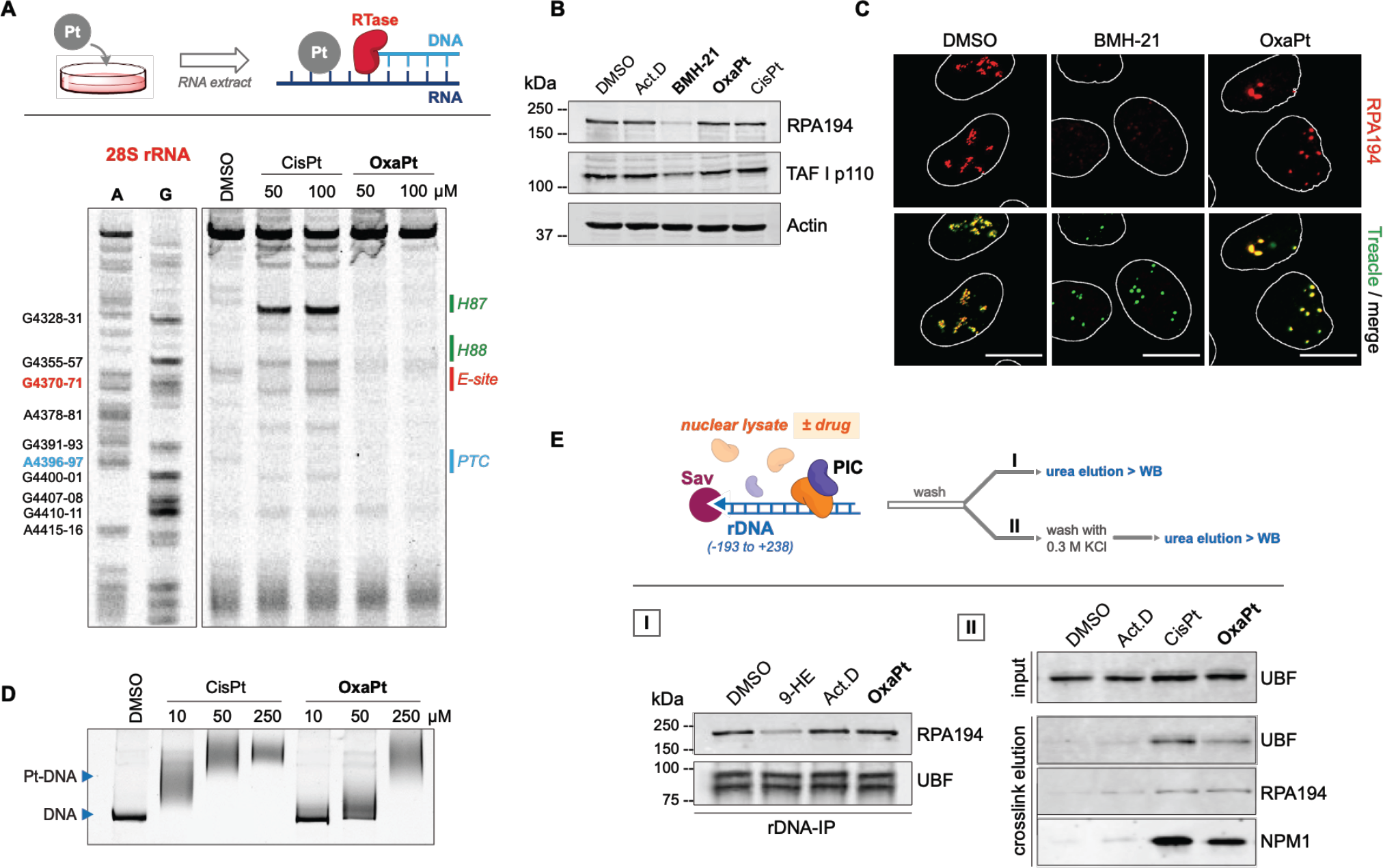
Oxaliplatin does not preferentially target nucleolar nucleic acids and transcription factors. (A) Schematic of primer extension assay for Pt adducts on cellular RNA *(top)*. Primer extension of a region in 28S human rRNA extracted from U2OS cells treated with 0-100 µM of cisplatin or oxaliplatin for 12 hours. Dideoxy sequencing ladder labeling A and G is shown on the left. Annotations of local rRNA structures are provided on the right. (B) Western blot of total cell extract from U2OS cells treated with Act. D (10 nM), BMH-21 (1 µM), oxaliplatin (10 µM), or cisplatin (10 µM) for 6 hours. Depletion of RPA194, and to a lesser extent TAF I p110, relative to actin control is seen only for BMH-21 treated cells. (C) Representative images of immunostaining against RPA194 (red) and Treacle (green) in cells treated as in (B). All scalebars = 15 µM. (D) Denaturing PAGE gel analysis of *in vitro* Pt binding to a 400-nt DNA fragment containing the ribosomal promoter sequence after incubation with cisplatin or oxaliplatin for 12 hours at 37°C in PtNA buffer (10 mM Na_2_PO_4_, 100 mM NaNO_3_, 1 mM Mg(NO_3_)_2_). Dose dependent labeling of the DNA is observed, with slower-migrating smeared bands indicating DNA with a greater number of Pt adducts. (E) Schematic of RNA Pol I pre-initiation complex (PIC) pulldown from extracted U2OS nuclear lysate using a streptavidin bead-immobilized DNA oligo containing the rDNA promoter sequence, with (I) or without (II) washing with 0.3 M KCl for detection of proteins covalently crosslinked to the DNA *(top)*. Western blot analysis of recovered large subunit of Pol I (RPA194) and UBF after incubation (1 hour, 4°C) of rDNA promoter with cell lysate containing DMSO (1% v/v), 9-hydroxyellipticine (40 µM), Actinomycin D (500 nM), or oxaliplatin (100 µM); depletion of RPA194 binding is seen only for 9-HE treatment *(bottom, left)*. Western blot analysis of recovered UBF, RPA194, and NPM1 after incubation of cell lysate with Act. D (500 nM), cisplatin (100 µM), or oxaliplatin (100 µM) for 1 hour at room temperature, followed by stringent washes containing 0.3 M KCl prior to elution into loading buffer *(bottom, right)*.

We next turned our attention to the rDNA promoter region, which is crucial for transcription initiation and has been identified as a target for small molecule inhibitors of Pol I (Ferreira et al., 2020). For example, treatment with the inhibitor BMH-21 has been shown to inhibit promoter escape and transcription elongation by Pol I leading to proteasome-mediated degradation of the Pol I large subunit (RPA194) (Jacobs et al., 2022). We did not observe depletion of RPA194 in oxaliplatin-treated cells, as measured by western blot (Fig. 3B) or immunofluorescence (Fig. 3C), indicating that oxaliplatin has a distinct mechanism of action from BMH-21. Instead, oxaliplatin treatment results in the partitioning of RPA194 into nucleolar cap structures, in agreement with previous studies of Pol I redistribution in response to transcriptional silencing (Shav-Tal et al., 2005).

Another potential mechanism of Pol I silencing is through inhibition of pre-initiation complex (PIC) assembly at the rDNA promoter (Andrews et al., 2013; Mars et al., 2020), so we investigated whether oxaliplatin is capable of perturbing these protein-DNA interactions. We first assayed the general reactivity of Pt compounds to DNA by treating a PCR amplified 400-nt rDNA segment containing the promoter sequence with various concentrations of Pt drug for 12 hr and analyzing adducts by denaturing PAGE. We observed substantially greater amounts of Pt-DNA adducts with 10-50 µM cisplatin as compared to 10-50 µM oxaliplatin (Fig. 3D), consistent with prior reports demonstrating greater inherent reactivity of cisplatin towards nucleic acids (Fichtinger-Schepman et al., 1985). Next, we performed pulldown assays to capture pre-initiation complex from U2OS nuclear extract using a biotinylated DNA fragment (Panov et al., 2001). When incubating immobilized rDNA promoter and nuclear extract in the presence of up to 100 µM oxaliplatin, we found that recruitment of Pol I and Upstream Binding Factor (UBF) to the promoter was unaffected (Fig. 3E, left). In contrast, we observed greatly reduced recovery of Pol I with the inhibitor 9-hydroxy-ellipticine, in agreement with its known ability to disrupt PIC assembly on the promoter (Andrews et al., 2013). To test the possibility that oxaliplatin may covalently crosslink rDNA and proteins, we adapted the method to add stringent washes with 0.3 M KCl after PIC capture, which has been shown to remove non-covalently bound PIC components. Under these conditions, we observed that 100 µM oxaliplatin treatment in nuclear extract resulted in the retention of Pol I, UBF, and NPM1 (Fig. 3E, right), indicative of covalent crosslinking. However, 100 µM cisplatin treatment resulted in similar or greater amounts of crosslinking with all three proteins, including UBF which has been previously reported to possess selective affinity for Pt-DNA adducts (Zhai et al., 1998). Given that cisplatin does not inhibit Pol I activity in cells at comparable concentrations to oxaliplatin, we conclude that interactions with the rDNA promoter and nucleolar proteins are unlikely to fully explain the specific effect of oxaliplatin on rRNA transcription. Further, covalent crosslinking with high concentrations of either Pt compound in nuclear extract does not mean that therapeutically relevant dosages of these compounds can crosslink rDNA and nucleolar proteins in cells.

Nevertheless, based on our finding that oxaliplatin can crosslink DNA and proteins in nuclear lysate, albeit at high concentrations, we investigated whether reactivity with nucleolar proteins could underlie transcriptional inhibition in cells. Recent work has shown that oxaliplatin can react with fibrillarin (FBL), an abundant scaffolding protein vital to nucleolar integrity, and proposed that crosslinking of FBL to nucleolar nucleic acids might be involved in oxaliplatin-mediated nucleolar disruption (Schmidt et al., 2021). To test this hypothesis, we transiently overexpressed FBL and measured how this impacted the activity of oxaliplatin, since overexpression was previously shown to confer resistance to oxaliplatin. While we observed a modest increase in cellular resistance to oxaliplatin upon FBL overexpression (SI Appendix, Fig. S3A), we saw no difference in nucleolar transcription compared to WT cells (SI Appendix, Fig. S3B). This result suggests that crosslinking of scaffolding proteins such as FBL by oxaliplatin, if it occurs, is unlikely to be a key driver for Pol I inhibition. In sum, our data indicates that transcriptional inhibition by oxaliplatin is not predominantly caused by direct interactions with biomolecules within the nucleolus, making its mechanism of action distinct from other major classes of Pol I inhibitors.

### Transcriptional silencing by oxaliplatin depends on ATM and ATR activity

Since Pt(II) drugs are known to activate the DDR in cells, we explored the possibility that oxaliplatin might inhibit Pol I transcription via a secondary response to Pt-DNA lesions occurring inside or outside of the nucleolus. Detection of genomic DSBs activates a wide range of signaling pathways throughout the cell, ranging from chromatin remodeling and proteasome recruitment to cell cycle arrest and programmed cell death (Blackford and Jackson, 2017). Notably, a number of studies have shown that global DNA damage or DNA damage targeted to the nucleolus can directly lead to transcriptional silencing (Korsholm et al., 2019; Kruhlak et al., 2007). Additionally, while transcriptional regulation at damaged genomic loci is a well-studied phenomenon, there is also evidence that DDR signaling can silence transcription distal to the DSB site (Shanbhag et al., 2010). In particular, Larsen *et al*. demonstrated that global silencing of nucleolar transcription can occur in response to DNA damage localized outside of the nucleolus by a mechanism involving ATM and other proteins associated with DSB recognition (Larsen et al., 2014). Although recent studies have questioned DDR activation as a major mediator of the effect of oxaliplatin on nucleolar transcription and organization (Bruno et al., 2017; Sutton and DeRose, 2021), the existence of specific DDR pathways communicating to nucleoli prompted us to investigate the relationship of oxaliplatin to DNA damage signaling.

To explore the connection between the DDR and nucleolar transcriptional inhibition, we used immunofluorescence microscopy to assay a panel of Pt compounds and Pol I inhibitors for the formation of γH2AX foci, a phosphorylated histone variant frequently used as a marker for DNA damage, in parallel with incorporation of 5-EU (SI Appendix, Fig. S4A). We observed robust induction of γH2AX foci within 6 hr of treatment with both cisplatin and oxaliplatin, though in agreement with previous reports, oxaliplatin-treated cells exhibited less γH2AX signal (SI Appendix, Fig. S4B). Interestingly, the Pol I inhibitor CX-5461 induced a greater number of γH2AX foci than oxaliplatin, consistent with a recent study reclassifying it as a DNA-damaging topoisomerase poison (Bruno et al., 2020). As expected, Act.D generated few γH2AX foci at the low dosage used for selective inhibition of Pol I (Choong et al., 2009). The induction of γH2AX foci by oxaliplatin motivated us to test whether DNA damage signaling precedes transcriptional inhibition by conducting a quantitative time-course analysis (SI Appendix, Fig. S4C). Although some foci are visible after 2 hr treatment with oxaliplatin, a general increase in γH2AX signal above untreated control is not seen until 3 hr, suggesting that extensive DNA damage signaling occurs after the onset of rRNA transcriptional silencing (SI Appendix, Fig. S4D). Based on these results, we conclude that although the extent of DNA damage induced by diverse agents does not appear to correlate strongly with Pol I inhibition, oxaliplatin is capable of activating DDR pathways concurrent with the induction of nucleolar stress.

Next, we explored the importance of upstream DDR signaling during oxaliplatin treatment. In canonical DDR signaling, recognition of DSBs is followed by activation of ATM and ATR kinases, which phosphorylate a wide range of targets to regulate DNA repair and cell cycle progression, among other cellular functions (Kinner et al., 2008). We investigated whether these pathways are relevant to oxaliplatin-mediated silencing of nucleolar transcription by co-treating cells with small molecule inhibitors for either ATM (KU-59933) or ATR (VE-821) and measuring the effect on 5-EU incorporation and nucleolar cap formation (Fig. 4A). Gratifyingly, inhibition of either ATM or ATR resulted in a partial rescue of transcription compared to oxaliplatin treatment alone, while inhibition of both kinases combined with oxaliplatin restored nucleolar transcription levels by nearly 70% (Fig. 4B). The formation of Treacle-containing nucleolar caps was also partially abrogated by these inhibitors, demonstrating that nucleolar segregation also relies on ATM and ATR activity. We next investigated whether the mechanism of other small molecule Pol I inhibitors depends on ATM/ATR, but we did not observe any transcriptional rescue by ATM/ATR inhibition for either Act.D or CX-5461 (Fig. 4C). Thus, oxaliplatin-induced silencing of rRNA transcription is mechanistically distinct from Act.D or CX-5461 and depends upon activation of ATM and ATR signaling. Further, we investigated whether downstream targets of ATM and ATR were involved in transcriptional silencing. We measured transcription levels in cells co-treated with oxaliplatin and inhibitors for the checkpoint kinases Chk1/Chk2, and once again observed rescue of nucleolar transcription (Fig. 4D), albeit to a lesser extent as compared to ATM/ATR inhibition. Taken together, these data indicate that oxaliplatin-induced rRNA silencing and nucleolar disruption occur downstream of the activation of ATM-Chk2 and ATR-Chk1 signaling pathways.

**Figure 4.**
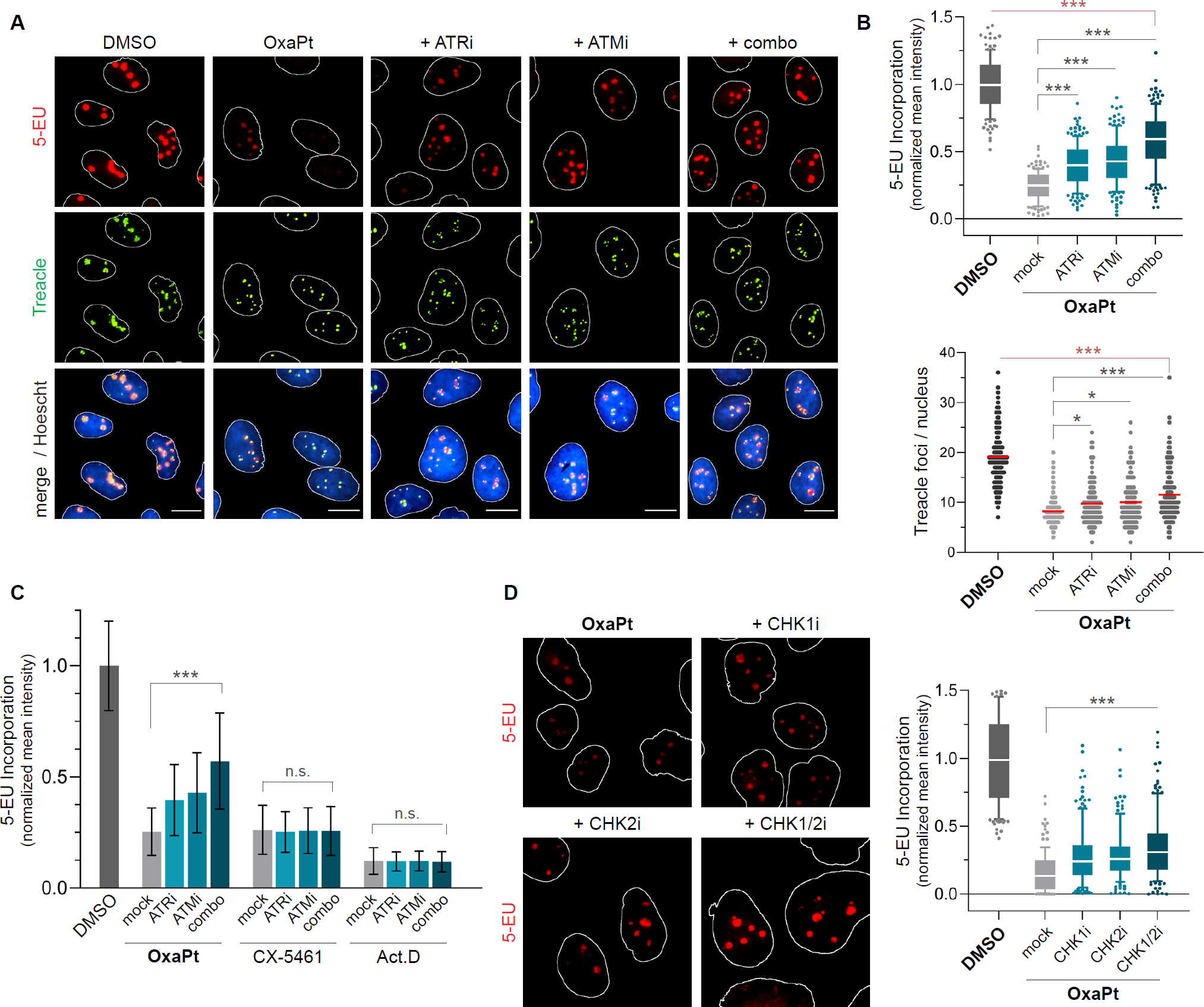
rRNA silencing by oxaliplatin is dependent on ATM and ATR activity. (A) Representative images of rRNA synthesis by 5-EU incorporation and localization of Treacle into nucleolar caps upon co-treatment of oxaliplatin (5 µM) with or without VE-821 (ATRi) and KU-55933 (ATMi) for 6 hours. (B) Quantification of 5-EU signal in nuclei from (A), normalized relative to DMSO control *(top)*. Quantification of Treacle foci count per nucleus from (A) *(bottom)*. Number of cells analyzed for each condition, summed over 3 biological replicates: DMSO (*N* = 141), oxaliplatin (*N* = 142), with ATRi (*N* = 143), with ATMi (*N* = 176), with both (*N* = 155). (C) Quantification of 5-EU signal in nuclei of cells co-treated with oxaliplatin (5 µM), CX-5471 (1 µM), or Actinomycin D (20 nM), with or without ATRi and ATMi, for 6 hours. At least *N* = 50 cells analyzed for each condition over 2 biological replicates. (D) Representative images of rRNA synthesis measured by 5-EU incorporation after co-treatment with oxaliplatin (5 µM) and inhibitors for CHK1 (300 nM), CHK2, (300 nM), or both CHK1/CHK2 (10 µM) *(left)*, and quantification of 5-EU incorporation, relative to DMSO control *(right)*. Number of cells analyzed in each condition, summed over 2 biological replicates: DMSO (*N* = 143), OxaPt/mock (*N* = 118), +CHK1i (*N* = 160), +CHK2i (*N* = 127), +CHK1/2i (*N* = 141). Statistical analysis in (B-D) was performed using a one-way analysis of variance, Tukey’s multiple comparisons test. Adjusted *p*-values are indicated by ***P<0.001, *P<0.05, and *n.s.* denotes a non-significant *p*-value > 0.05. All experiments were performed in wild-type U2OS cells. All scalebars = 15 µM.

### The nucleolar response to oxaliplatin shares features with canonical n-DDR signaling

We next sought to characterize the mechanism that bridges DNA damage signaling to nucleolar reorganization and transcriptional inhibition by oxaliplatin. Based on our observation that Treacle colocalizes with oxaliplatin-induced nucleolar caps, we considered the possibility that a nucleolar DNA damage response (n-DDR) is operative in this context. Previous studies of n-DDR have shown that DSB induction by ionizing radiation (IR) or I-Ppo1, a homing endonuclease which selectively cleaves sites in rDNA, results in transcriptional silencing by a mechanism centered around Treacle-mediated recruitment of DDR factors to nucleoli (Kruhlak et al., 2007; van Sluis and McStay, 2015). In particular, Treacle has been proposed to interact directly with the DDR adapter protein NBS1, a critical component of the MRE11-RAD50-NBS1 (MRN) complex involved in DSB repair (Ciccia et al., 2014; Larsen et al., 2014). Using an antibody against endogenous NBS1, we found that 31% of oxaliplatin-treated cells contain NBS1 perinucleolar foci that co-localize with UBF-positive nucleolar caps (Fig. 5A). We performed a time-course study to track NBS1 localization at various stages of inhibition by oxaliplatin (SI Appendix, Fig. S5) and observed recruitment of NBS1 to the interior of nucleoli at the onset of inhibition (∼2-3 hr) prior to segregation into peripheral cap structures at later time points, in agreement with a previous report using I-Ppo1 (Mooser et al., 2020). We next asked if oxaliplatin-induced NBS1 recruitment to nucleoli is dependent on Treacle. Upon knockdown of Treacle by RNAi, we observed nearly complete abrogation of nucleolar recruitment of NBS1 in both oxaliplatin-treated and I-Ppo1 transfected cells (Fig. 5B), suggesting that the Treacle-NBS1 interaction is operative in both conditions. Notably, we found that NBS1 recruitment is not a general feature of Pol I inhibition, as fewer than 5% of Act.D and doxorubicin-treated cells and only 15% of CX-5461 treated cells contained NBS1-positive caps after nucleolar segregation (Fig. 5C). In contrast, transfection with I-Ppo1 resulted in nucleolar NBS1 recruitment in 87% of transfected cells. Given that doxorubicin and CX-5461 are potent DNA damaging agents, these results indicate that Treacle-mediated recruitment of NBS1 is specific to certain forms of DNA damage that activate the n-DDR response.

**Figure 5.**
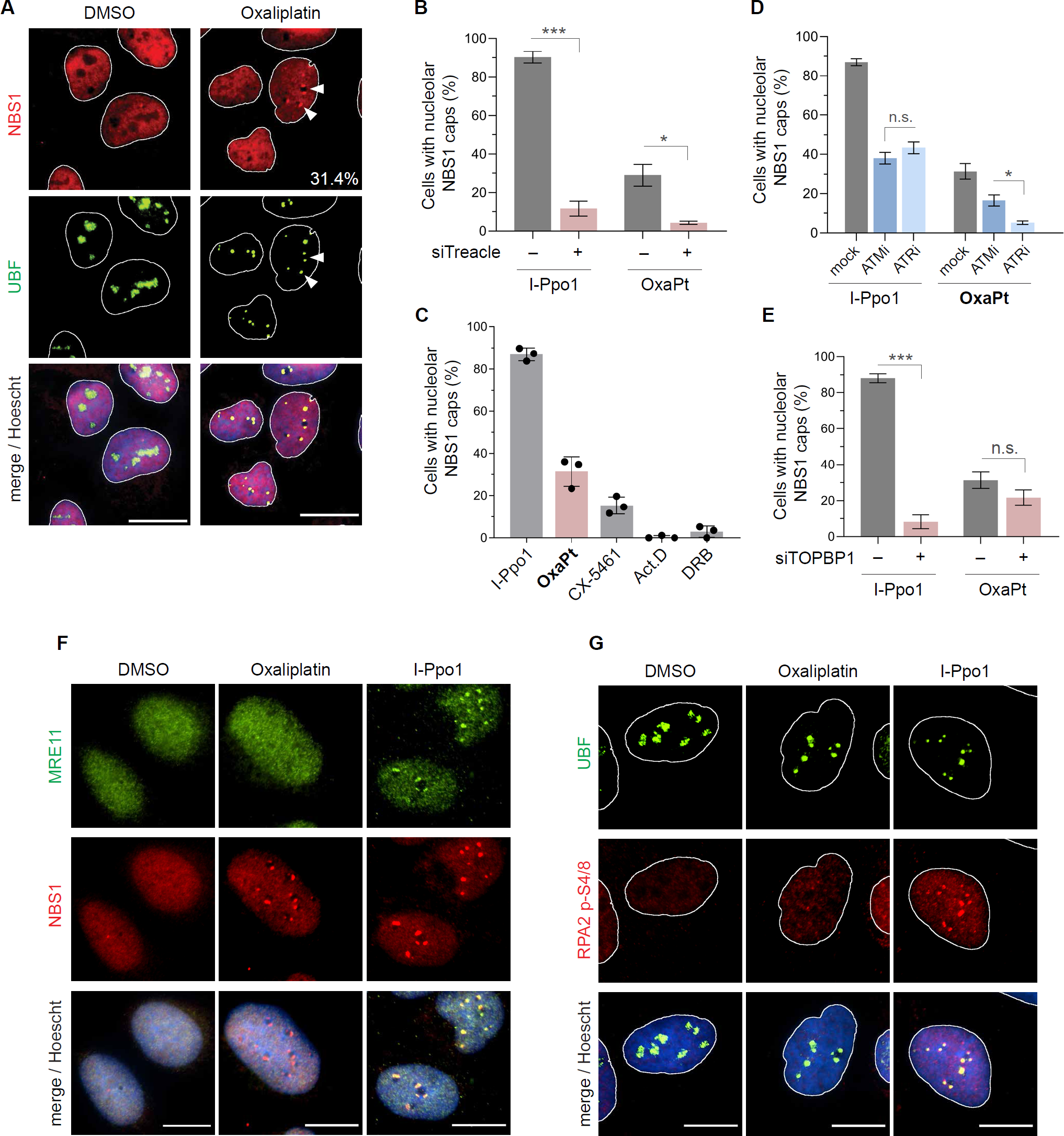
Oxaliplatin induces nucleolar recruitment of NBS1 but not of other repair-associated DDR factors. (A) Representative images of immunostaining against NBS1 (red) and UBF (green) in cells treated with or without oxaliplatin. White arrows point to instances of NBS1 colocalization with UBF-positive nucleolar caps. Number in bottom right indicates percentage of cells exhibiting nucleolar caps containing NBS1. All scalebars = 20 µM. (B) Fraction of I-Ppo1 or oxaliplatin-treated cells containing NBS1-positive nucleolar caps with or without siRNA knockdown of endogenous Treacle. (C-D) Quantification of the fraction of cells containing NBS1-positive nucleolar caps out of all cells with nucleolar caps, as indicated by peripheral UBF foci around nucleoli, after treatment with indicated drugs or I-Ppo1. Error bars represent mean ± SEM of 3 biological replicates. (E) Fraction of I-Ppo1 or oxaliplatin-treated cells containing NBS1-positive nucleolar caps with or without siRNA knockdown of endogenous TOPBP1. Statistical analysis in (B-E) was performed using a one-way analysis of variance, Tukey’s multiple comparisons test. Adjusted *p*-values are indicated by ***P<0.001, *P<0.05, and *n.s.* denotes a non-significant *p*-value > 0.05. (F-G) Representative images of immunoblotting against MRE11 and NBS (F) or UBF and phospho-RPA2 (G) after treatment with I-Ppo1 or oxaliplatin. All experiments were performed in wild-type U2OS cells. All scalebars = 20 µM.

Nucleolar segregation and NBS1 recruitment by Treacle in the context of n-DDR have been shown to depend on ATM and ATR signaling (Ciccia et al., 2014; Larsen et al., 2014). As demonstration of this, we pre-treated cells with ATM or ATR inhibitors prior to transfection by I-Ppo1 and observed substantially fewer NBS1-positive nucleolar caps, as previously reported (Fig. 5D). When we combined either kinase inhibitor with oxaliplatin treatment, we again saw a reduction in NBS1 recruitment to nucleoli. However, in the case of oxaliplatin this phenomenon was more sensitive to ATR inhibition, as only 5% of ATR-inhibited cells showed NBS1-positive nucleolar caps compared to 17% of ATM-inhibited cells, while NBS1 recruitment by I-Ppo1 was equally suppressed in both cases. A previous study of the n-DDR response to I-Ppo1 showed that NBS1 recruitment is also dependent on TOPBP1, an ATR activator proposed to interact directly with nucleolar Treacle (Mooser et al., 2020). Although we observed that TOPBP1 knockdown by RNAi indeed resulted in the inhibition of NBS1 recruitment in I-Ppo1 transfected cells, we were surprised to see no change in NBS1 recruitment by oxaliplatin under these conditions (Fig. 5E). Interestingly, we still detected colocalization of TOPBP1 with nucleolar caps in a subset of oxaliplatin-treated cells, despite its apparent lack of interaction with NBS1 in this context (SI Appendix, Fig. S6).

Given that we only observed NBS1 and TOPBP1 in a fraction of oxaliplatin-treated cells, we wanted to investigate whether their role is linked to cell cycle state. Both NBS1 and TOPBP1 have been shown to be essential modulators of cell cycle checkpoint control through their interactions with ATM and ATR (Wardlaw et al., 2014; Zhang et al., 2006). Since the checkpoint response to DNA damage is typically activated by replication stress (Panagopoulos and Altmeyer, 2021), we established that γH2AX activation by oxaliplatin occurs almost exclusively in S-phase cells by EdU incorporation (SI Appendix, Fig. S7A). We further found strong cell cycle dependency of nucleolar NBS1 recruitment, with ∼60% of EdU-positive cells exhibiting nucleolar NBS1 foci as compared with less than 10% of EdU-negative cells (SI Appendix, Fig. S7B). However, even in non-S-phase cells lacking observable γH2AX foci and nucleolar NBS1, we still observed signs of nucleolar disruption. Based on these results, we reason that oxaliplatin activates DDR pathways involving NBS1 in cells undergoing DNA replication, but these responses are unlikely to be the universal mechanism of nucleolar stress induction by oxaliplatin in all cells. Thus, the nucleolar response to oxaliplatin appears to be distinct from the canonical n-DDR mechanisms for rDNA damage, despite sharing a few key similarities.

### NBS1 and TOPBP1 are not essential for oxaliplatin-mediated rRNA synthesis inhibition

Our finding that the nucleolar response to oxaliplatin differs from that of I-Ppo1 prompted us to explore similarities to an alternative n-DDR pathway which responds to DSBs outside of the nucleolus. As reported by Larsen *et al*. (Larsen et al., 2014), this *in-trans* n-DDR mechanism is distinguished by lack of MRE11 recruitment to nucleoli, indicating the absence of active DNA repair at the rDNA loci sequestered in nucleolar caps. Moreover, many individual nucleoli silenced by this pathway also lack γH2AX accumulation, in contrast to the result of direct rDNA damage targeted to nucleoli (Mooser et al., 2020; van Sluis and McStay, 2015). To investigate the possibility that oxaliplatin activates an *in-trans* DNA damage response, we first checked for the presence of γH2AX foci in the vicinity of nucleoli. While most oxaliplatin-induced γH2AX foci were distributed throughout the nucleoplasm, a minor fraction of cells exhibited faint γH2AX foci overlapping with Treacle-positive nucleolar caps (SI Appendix, Fig. S7C, top). However, these foci were considerably less intense and frequent than those observed for I-Ppo1 transfected cells (SI Appendix, Fig. S7C, bottom), suggesting that oxaliplatin induces a much weaker DNA damage response within nucleoli compared to I-Ppo1, if at all. Furthermore, we did not observe any colocalization of endogenous MRE11 with NBS1 foci in oxaliplatin-treated cells (Fig. 5F), indicating that NBS1 is operating independently of the MRN repair complex. We also saw no phosphorylated RPA2 (RPA2 pS4/S8) in NBS1-containing nucleolar caps (Fig. 5G), suggesting that oxaliplatin does not induce active DNA end resection within these structures. In contrast, transfection of I-Ppo1 resulted in the association of both MRE11 and p-RPA2 with nucleolar caps, consistent with the mechanism for *in-cis* n-DDR signaling. Based on the absence of these repair factors and minimal γH2AX signaling within nucleoli, we conclude that oxaliplatin activates some form of *in-trans* DDR signaling to the nucleolus.

Although our earlier results indicated that nucleolar recruitment of NBS1 is not a general feature of oxaliplatin-induced nucleolar stress, all reported n-DDR mechanisms to date have proposed a role for NBS1 in rRNA silencing. To further explore the importance of NBS1 for transcriptional inhibition by oxaliplatin, we utilized a U2OS cell line expressing a hypomorphic C-terminal fragment of NBS1, hereafter referred to as “NBS1ΔN”. Since the N-terminus of NBS1 is necessary for interactions with Treacle and the repair factor CtIP, among other DDR proteins, this cell line is deficient in DNA end resection and does not undergo nucleolar segregation or rRNA silencing in response to rDNA damage by I-Ppo1 (Anand et al., 2019; Mooser et al., 2020). When we treated NBS1ΔN cells with oxaliplatin, we were surprised to observe an even greater decrease in rRNA transcription relative to wild-type U2OS cells (Fig. 6A, 6B), while transcription during I-Ppo1 transfection remained much higher in NBS1ΔN cells as previously reported. However, oxaliplatin-induced transcriptional inhibition was still partially reversed in NBS1ΔN cells by co-treatment of inhibitors for ATM and ATR (Fig. 6C), suggesting that ATM/ATR-dependent transcriptional silencing remains operative even in the absence of repair-competent NBS1. Furthermore, we did not observe any difference in nucleolar transcription for oxaliplatin-treated cells lacking TOPBP1, whereas I-Ppo1 induced silencing was substantially reversed under these conditions (SI Appendix, Fig. S8A, S8B). Together, these results demonstrate that although nucleolar recruitment of NBS1 and TOPBP1 is observed in a subset of oxaliplatin-treated cells, neither is essential for ATM/ATR signaling to nucleoli and subsequent transcriptional inhibition, in contrast to reported n-DDR mechanisms. Nonetheless, we determined that NBS1ΔN cells are more than 2-fold more sensitive to oxaliplatin compared to wild-type cells, likely due to widespread DNA repair defects at oxaliplatin-induced damage sites (Fig. 6D). By comparison, sensitization of NBS1ΔN cells to cisplatin was observed to a lesser extent, suggesting that NBS1 plays an important role in the repair of oxaliplatin-induced lesions but does not mediate transcriptional inhibition and nucleolar disruption. Based on these findings, we propose that oxaliplatin inhibits rRNA transcription through activation of NBS1-independent *in-trans* n-DDR signaling in response to DNA damage throughout the nucleus.

**Figure 6.**
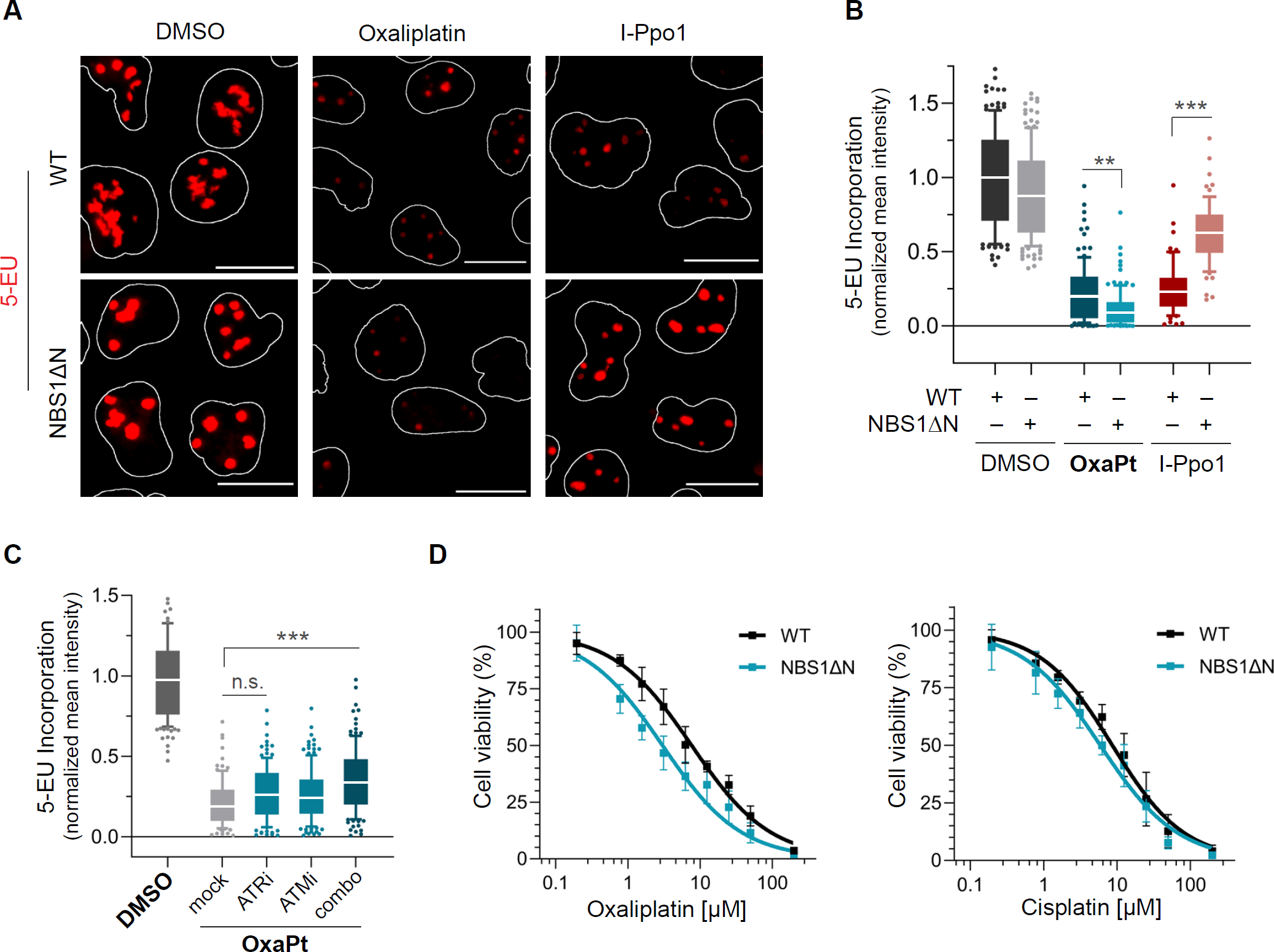
Full NBS1 function is not essential for rRNA silencing by oxaliplatin. (A) Representative images of rRNA synthesis in wild-type and “NBS1ΔN” U2OS cells as measured by 5-EU incorporation after treatment with oxaliplatin (5 µM, 6 hours) or transfection of I-Ppo1 (8 hours). All scalebars = 15 µM. (B) Quantification of 5-EU signal from (A), relative to DMSO control in WT cells. Number of cells analyzed in each condition, summed over 2 biological replicates: DMSO in WT (*N* = 143) and NBS1ΔN (*N* = 142), oxaliplatin in WT (*N* = 112) and NBS1ΔN (*N* = 116), and I-Ppo1 in WT (*N* = 67) and NBS1ΔN (*N* = 65). For I-Ppo1-treated cells, only cells positive for transfection by HA staining were quantified. (C) Quantification of 5-EU signal in cells co-treated with oxaliplatin and ATRi, ATMi, or both, relative to DMSO control. At least *N* = 100 cells analyzed for each condition over 2 biological replicates. Statistical analysis in (B-C) was performed using a one-way analysis of variance, Tukey’s multiple comparisons test. Adjusted p-value is indicated by ***P<0.001, **P<0.01, and *n.s.* denotes a non-significant *p*-value > 0.05. (D) Dose response curves for oxaliplatin and cisplatin in wild-type and NBS1ΔN U2OS cells. Cell viability was measured using MTS assay at 72 hours post-treatment and corrected for DMSO control in each cell line. Error bars represent mean ± SEM of *N* = 6 biological replicates.

## DISCUSSION

In this manuscript, we investigate the cellular signaling pathways responsible for coordinating the disruption of nucleolar organization and rRNA transcription by the clinically used Pt-drug oxaliplatin. We show that oxaliplatin-induced rRNA silencing and nucleolar disruption occur downstream of the ATM-Chk2 and ATR-Chk1 pathways, and we demonstrate that small-molecule inhibition of either pathway restores nucleolar transcription and structure. Further, we find that oxaliplatin does not induce substantial amounts of nucleolar DNA damage, and that oxaliplatin-induced rRNA silencing does not depend upon NBS1 or TOPBP1, distinguishing it from previously characterized *cis* and *trans* n-DDR pathways. Our findings indicate that oxaliplatin induces a distinct DDR signaling pathway that functions *in trans* to inhibit Pol I transcription and induce nucleolar stress, resulting in activation of p53 and disruption of ribosome biogenesis, and ultimately cell death. Our study demonstrates a previously underappreciated connection between DNA damage signaling and nucleolar stress, and illuminates the diverse mechanisms of action exhibited by chemotherapeutic Pt drugs.

By studying the nucleolar accumulation and function of specific DDR factors, we demonstrate that nucleolar stress induction by oxaliplatin shares many characteristics with previously described nucleolar DDR signaling pathways. In particular, the lack of MRE11 and phospho-RPA2 in nucleoli is consistent with previously reported *in-trans* n-DDR signaling resulting from DNA damage targeted outside of nucleoli (Larsen et al., 2014), and distinguishes oxaliplatin-induced DDR signaling from *in-cis* n-DDR pathways that respond to and repair targeted rDNA damage. Notably, while we observe Treacle-dependent NBS1 and TOPBP1 foci within nucleolar cap structures, neither protein is required for oxaliplatin-induced rRNA silencing, in contrast to prior characterization of *trans* n-DDR signaling. Since both NBS1/TOPBP1 accumulation and transcriptional silencing depend upon ATM/ATR activity, we posit that these pathways function in parallel upon oxaliplatin induction of DDR signaling. The precise mechanism of Pol I inhibition through ATM/ATR signaling is not yet clear. Some Pol I transcription factors are known targets of ATM and ATR (Matsuoka et al., 2007), while ATM-dependent post-translational modification of histones has been shown to regulate rDNA transcription in response to IR-induced DNA damage (Pefani et al., 2018). Given the apparent mechanistic diversity in nucleolar DDR signaling, further studies are necessary to characterize nucleolar ATM/ATR substrates in response to different forms of DNA damage.

Our finding that NBS1 is recruited to nucleoli independently of MRE11 by oxaliplatin is notable, as little is known about the cellular functions of NBS1 outside of the MRN repair complex. In oxaliplatin-treated cells, the appearance of nucleolar NBS1 correlates strongly with the induction of replication stress via γH2AX signaling in S-phase, suggesting there may be a threshold of DNA damage required for its recruitment to nucleoli. Although the absence of MRE11 and phospho-RPA2 suggests otherwise, we cannot rule out that a small amount of DNA repair occurs in nucleolar caps containing NBS1. However, given that we observe comparable rRNA silencing in cells lacking nucleolar NBS1 and γH2AX foci, we conclude that transcriptional silencing in this context likely occurs via an NBS1-independent pathway. Nevertheless, we show that nucleolar NBS1 recruitment is not seen upon transcriptional inhibition of Pol I by other DNA-damaging drugs like CX-5461 and doxorubicin, so its presence in the cellular response to oxaliplatin is noteworthy and warrants further investigation.

The cytotoxicity of Pt drugs is generally thought to be driven by their ability to form Pt-DNA adducts, yet our report and previous studies have shown that oxaliplatin generates fewer DNA crosslinks per base than cisplatin, but is comparably cytotoxic (Woynarowski et al., 2000). While these observations have led researchers to speculate that oxaliplatin-mediated nucleolar stress and associated transcription/translation inhibition occurs through a DNA-damage independent mechanism, our study shows that DDR signaling is central to oxaliplatin-mediated nucleolar disruption and rRNA silencing. In contrast, cisplatin fails to induce nucleolar phenotypes except at artificially high concentrations well above clinically relevant dosages. How do these two structurally related Pt drugs generate dramatically different cellular outcomes? We propose that DDR signaling in response to oxaliplatin-DNA lesions is driven by specific recognition of oxaliplatin-DNA adducts rather than by the total amount of generated DNA damage. In support of this idea, biophysical analyses of cisplatin and oxaliplatin adducts on DNA have identified differences in bending angle, hydrogen bonding patterns, and conformational flexibility of the local DNA structure, all of which have been proposed to affect the affinity of damage recognition proteins for Pt adducts (Jamieson and Lippard, 1999; Malina et al., 2007; Spingler et al., 2001). In addition, several HMG-domain proteins, such as UBF and HMGB1, as well as mismatch repair protein MSH2, have been demonstrated *in vitro* to bind cisplatin-DNA and oxaliplatin-DNA adducts with different affinities, which may result in differential recognition and DDR signaling in cells (Wei et al., 2001; Zdraveski et al., 2002; Zhai et al., 1998). Based on this reasoning, we speculate that the size and shape of oxaliplatin lesions lead to distinct recognition by DDR factors coupled to nucleolar signaling.

An alternative hypothesis to explain the unique ability of oxaliplatin among Pt drugs to induce nucleolar stress is through partitioning and enrichment in the nucleolar compartment. Interestingly, recent work has highlighted the propensity for some antineoplastic drugs to preferentially partition into phase-separated molecular condensates including the nucleolus (Klein et al., 2020). It is currently unknown whether this behavior is driven by specific small molecule-condensate interactions or through general physicochemical properties such as molecular size and hydrophobicity, however the increased hydrophobicity of the oxaliplatin DACH ligand in comparison to polar cisplatin NH_3_ ligands makes a partitioning mechanism worthy of further exploration. While our manuscript was in preparation, Schmidt *et al*. reported that oxaliplatin can crosslink rRNA to the nucleolar scaffolding protein FBL, and suggested that oxaliplatin may selectively accumulate in nucleoli (Schmidt et al., 2021). While we cannot formally exclude this possibility, we do not favor this hypothesis based upon several data points presented herein. First, we show by RT-primer extension that cisplatin accumulates to a greater degree than oxaliplatin on 28S, 18S, and 5.8S rRNA molecules. Second, we do not observe significant DNA damage foci in the nucleolus or robust activation of *in-cis* n-DDR after oxaliplatin treatment as would be expected if the compound were to selectively partition in this compartment. Finally, overexpression of FBL does not lead to restoration of nucleolar structure or rRNA synthesis. In addition, a recent structure-activity study of oxaliplatin-like compounds by McDevitt *et al*. demonstrated that hydrophobicity alone did not correlate with nucleolar stress induction, and instead identified the existence of subtle structural requirements on the Pt diamine ligand (McDevitt et al., 2022). Taken together, these results favor a model where specific recognition of oxaliplatin-DNA lesions occurring throughout the nucleus (rather than specifically within the nucleolus) is the major driver for nucleolar stress and cytotoxicity. Further characterization of these lesions in the context of nucleolar signaling will be necessary to identify the DDR factors responsible for specific recognition. As highlighted in this study, the disruption of ribosome biogenesis through specific DNA damage signaling pathways is a promising therapeutic strategy and may inform development of the next generation of Pol I transcription inhibitors.

## SIGNIFICANCE

Pt-based drugs are an important class of anti-cancer therapeutics in widespread clinical use, but a number of key questions remain relating to their mechanism of action. Historically, Pt compounds have been proposed to function as DNA damaging agents, but recent studies have demonstrated that oxaliplatin, one of the three FDA-approved Pt drugs, is better characterized as a transcription/translation inhibitor. These findings suggest that oxaliplatin kills cells in a fundamentally different manner from other Pt drugs, however the underlying molecular mechanism remains unknown. In this work, we characterize a novel pathway that links DNA damage signaling to transcriptional inhibition in the nucleolus by oxaliplatin. We show that selective silencing of rRNA transcription in this context is mediated by the DNA damage response kinases ATM and ATR. Further, we find that oxaliplatin does not target nucleolar proteins or nucleic acids, and rather that this DNA damage signal likely originates from oxaliplatin-DNA lesions occurring outside of the nucleolus. We draw parallels to a reported *in-trans* nucleolar DNA damage response pathway, but ultimately show that the response to oxaliplatin is mechanistically distinct. In this manner, our study illuminates the mechanistic diversity of nucleolar DDR signaling. Moreover, we compare oxaliplatin against other small molecule inhibitors of rRNA synthesis and find that oxaliplatin is unique in its reliance upon DNA damage signaling. We anticipate that this mechanistic discovery will expand the repertoire of biological pathways that can be targeted by further development of therapeutic inhibitory agents.

## ACKNOWLEDGMENTS

We thank Joshua Riback for helpful discussions regarding nucleolar organization. We are grateful to Manuel Stucki for providing the NBS1ΔN cell line. R.E.K. acknowledges support from the National Institute of Health (R01 GM132189), the Alfred P. Sloan Foundation, and the Princeton University Innovation Fund for New Ideas in the Natural Sciences. M.N. was supported by a generous gift from the Edward C. Taylor 3rd Year Graduate Fellowship in Chemistry. All authors acknowledge financial support from Princeton University.

## AUTHOR CONTRIBUTIONS

M.N. conducted experiments and wrote the paper. R.E.K. supervised M.N. and wrote the paper.

## COMPETING INTEREST STATEMENT

The authors declare no competing financial interests.

## Notes

### Competing Interest Statement

The authors have declared no competing interest.

### Summary of Updates

Revised on 04/03/2023.

