## Supplementary Information for "Oxaliplatin Inhibits RNA Polymerase I via DNA Damage Signaling Targeted to the Nucleolus"

Table of contents

| | |
|---|---|
| Materials and Methods | 2 |
| Supplementary Tables S1-S3 | 6 |
| Supplementary Figures S1-S8 | 9 |

### Materials and Methods

**Compounds.** 5-ethynyl uridine (5-EU) and 2'-azidocytidine (2'-AzCyd) were purchased from Carbosynth. THPTA, Cy3-alkyne, and Cy3-DBCO were obtained from Click Chemistry Tools. CX-5461, BMH-21, doxorubicin, 9-hydroxy-ellipticine, GDC-0575 (CHK1i), AZD7762 (CHK1/2i), and BML-277 (CHK2i) were purchased from Cayman Chemical. All other chemicals were purchased from Sigma-Aldrich or Fisher Scientific unless otherwise indicated. Stock solutions of compounds were made as follows: 5 mM stocks of cisplatin and phenanthriplatin were prepared in DMF; 5 mM stocks of oxaliplatin were prepared in water; 5 mM stocks of VE-821, KU-55933, 9-hydroxy-ellipticine, and 2 mM stocks of actinomycin D, CX-5461, BMH-21, and CHK1/2 inhibitors were prepared in DMSO. All platinum drug stocks were prepared fresh weekly and aliquoted for storage at -20°C. Stock solutions were diluted into DMEM media immediately prior to treatment of cell cultures.

**Cell Culture and Drug Treatments.** HeLa (cervical carcinoma cell line) and U2OS (human osteosarcoma cell line) cells were cultured at 37 °C in a humidified atmosphere with 5% $CO_2$ in DMEM (Thermo Fisher, 11995073) supplemented with 10% fetal bovine serum (Bio-Techne, S12450H), 2 mM L-glutamine (Thermo Fisher, 25030–081), and penicillin-streptomycin antibiotics (Thermo Fisher, 15070-063). U2OS cells expressing the "NBS1ΔN" truncated protein were obtained as a kind gift from Manuel Stucki (University of Zurich) (1). All drug treatments were conducted on cells grown to 70% confluency.

Except where noted otherwise, cisplatin and oxaliplatin drug treatments were carried out at a final concentration of 5 μM Pt drug for 6 hours. For pulsed 5-EU incorporation to visualize nucleolar transcription, cells were incubated for 1 hour in medium containing 1 mM 5-EU. To measure transcription during drug treatment, medium containing the drug would be replaced with fresh medium containing both 1 mM 5-EU and the drug compound for the final 1 hour of the treatment. To analyze cell cycle state, cells were incubated for 1 hour in medium containing 10 μM 5-EdU at the end of the corresponding drug treatment. For experiments involving co-treatment with a kinase inhibitor, cells were pre-treated with inhibitor alone for 4 hours prior to addition of fresh media containing both compounds. Unless stated otherwise, the following compounds were used at the indicated final concentrations: ATR inhibitor VE-821 (4 μM), ATM inhibitor KU-55933 (10 μM), CHK1 inhibitor GDC-0575 (300 nM), CHK1/CHK2 inhibitor AZD-7762 (300 nM), and CHK2 inhibitor BML-277 (10 μM).

**Plasmids and RNA Transfection.** The construct for NPM1-mCherry expression was generated by PCR amplification from FM5-NPM1-mCherry (gift from Clifford Brangwynne, Princeton University) and ligation into a modified pcDNA5/FRT/TO (Life Technologies, V6520-20) vector containing an N-terminal 3xFLAG tag. HA-NLS-I-PpoI cDNA was obtained from Addgene (46963). For transient transfection, HA-NLS-I-PpoI cDNA was cloned into pcDNA5/FRT/TO vector (Life Technologies, V6520–20). DNA plasmids were transfected into cells with Lipofectamine 2000 (Invitrogen) according to the manufacturer's instructions. Immunofluorescence analysis of I-Ppo1 was performed 8 hours after the transfection. For silencing of protein expression by RNAi, the control siRNA (siCtrl) and siRNA against TOPBP1 (siTOPBP1) and Treacle (siTCOF) were obtained from Dharmacon (Horizon Discovery). The sequences of siRNAs used in this study are listed in Table S2. Cells were grown in six-well plates 24 hours prior to the transfection. The siRNA transfection was done with 20 pM siRNA using Lipofectamine RNAiMAX (Invitrogen). Cells were split on coverslips 24 hours after the transfection and used for immunofluorescence 72 hours after transfection.

**Immunofluorescence.** Cells to be imaged were grown on 12 mm coverslips (Fisher Scientific, 12-545-81) in 24-well plates as described above. After treatment, cells were washed once with ice-cold PBS, fixed for 20 minutes at RT with PBS containing 3% paraformaldehyde adjusted to pH 7.3, and then washed 3 times for 5 minutes each with 1×PBS. Next, cells were permeabilized with 0.5% Triton-X in PBS for 20 minutes at RT. For click chemistry labeling of cellular RNA in 5-EU and 5-EdU treated samples, coverslips were incubated in 100 μL drops of freshly prepared CuAAC reaction mixture containing Cy3-alkyne (10 μM, Click Chemistry Tools, TA117-1), $CuSO_4$ (1 mM), THPTA ligand (2 mM), and sodium ascorbate (10 mM) in PBS. The reaction was allowed to proceed for 2 hours at room temperature in the dark. Cells were washed three times in PBST (0.1% Triton-X100 in 1×PBS) for 15 minutes each to remove free Cy3 dye. For effective visualization of nucleolar Cy3 signal for 5-EU treated cellular RNA, further immunostaining or incubation in blocking buffer was necessary to wash out nucleoplasmic signal. For immunostaining, the coverslips were blocked with 5% goat serum or 3% BSA in PBST for 1 hour, then incubated with primary antibody for 2 hours at room temperature. After washing three times with PBST for 5 minutes each, secondary antibody incubations were performed for 1 hour at RT in the dark. All antibodies used in this study are listed in Table S1. The coverslips were then washed twice more with PBST for 5 minutes each, stained with Hoescht 33342 (Thermo Scientific, 1 ug/mL) for 10 minutes, washed with PBS once more, and mounted on glass microscopy slides in ProLong AntiFade Reagent (Life Technologies) and sealed with nail polish.

**Imaging and Quantification.** Images of fixed cells were acquired using NIS Elements AR software and a Nikon Eclipse Ti microscope equipped with 100x objective and CMOS camera. Images used for direct comparison were acquired using standardized illumination and exposure settings and displayed with identical lookup table (LUT) settings. For optimal representation in figures, images were adjusted for brightness and exported as RBG TIF files using ImageJ. Quantification of immunofluorescent signal was done in ImageJ. Segmentation of nuclei was performed by generating regions of interest (ROI) from the DAPI channel. The mean fluorescence intensity of nucleolar 5-EU was approximated by the calculated intensity across nuclear ROIs in the EU channel, which contained minimal nucleoplasmic signal due to the short-pulse labeling and optimized staining conditions. Quantification of NPM1 localization was performed by measuring the CV for individual nuclei, defined as the standard deviation in pixel intensity in ROIs divided by the mean pixel intensity, which was normalized relative to the untreated control.

**In-gel Fluorescence.** For gel electrophoresis analysis of ongoing rRNA transcription, U2OS WT cells were pre-treated with indicated drug compound for 2 hours, then co-treated with drug and 1 mM 2'-azidocytidine for an additional 6 hours. Cells were harvested by scraping in 1×PBS and total RNA was extracted using TRIzol LS (Invitrogen) according to the manufacturer's instructions. RNA samples were then reacted with Cy3 fluorophore using SPAAC conditions; 15 µg of total RNA was combined with 100 µM Cy3-DBCO (Click Chemistry Tools, A140-1) and 1X PBS in a 20 µL reaction volume, containing 0.5 µL of murine RNase inhibitor (NEB, M0314), and incubated for 2 hours at 37 °C. All reactions were purified using Zymo RNA Clean and Concentrator-5 spin columns according to the manufacturer's instructions and separated by native gel electrophoresis using 1% TAE-agarose. Labeled RNA was visualized by in-gel fluorescence on a Typhoon FLA 9500 Fluorescent Image Analyzer Scanner (GE Healthcare) using a Cy3 filter set. Total RNA was visualized by staining with ethidium bromide.

**rDNA Promoter Pulldown of Nucleolar Proteins.** The biotinylated rDNA promoter (-193 to +238) was synthesized by PCR amplification using the plasmid prHu3 (gift from Robert Tjian, UC Berkeley) as a template (2). The primers used are listed in Table S3; the sense primer was biotinylated at the 5' end. The resulting 5'-end-biotinylated DNA fragments were gel purified and immobilized on streptavidin-coated paramagnetic beads (M280 Dynabeads, Invitrogen) according to the manufacturer's instructions. In a typical reaction (3), 750 ng of DNA was immobilized on 15 uL of beads, blocked with 5% BSA, then incubated with gentle agitation in U2OS cell nuclear extract for 30 minutes at 4°C in 50 mM KCl – TM10i buffer (50 mM Tris HCl pH 7.4, 12.5 mM $MgCl_2$, 1 mM EDTA, 10% glycerol, 1 mM DTT, 0.03% NP-40) containing EDTA-free protease inhibitor cocktail (Roche). After separation using a magnetic stand, beads were washed three times with 2 reaction volumes of TM10i-0.05M KCl buffer. Captured proteins were eluted by boiling in SDS loading buffer and analyzed by immunoblotting.

**Primer Extension Analysis.** For primer extension analysis of Pt adducts on cellular RNA, HeLa cells were treated with the indicated Pt compounds for 12 hours, then harvested by scraping into ice-cold 1×PBS solution. Total RNA was extracted using TRIzol LS (Invitrogen) according to the manufacturer's instructions. After DNase I treatment (NEB, M0303) and purification using Zymo RNA Clean and Concentrator-5 columns, 4 µg of total RNA was annealed to 10 pmol of IR-labeled primer complementary to various segments of rRNA (listed in Table S3) by heating to 65°C for 3 minutes and slow cooling (0.5°C/sec) to 4°C. Primer extension was performed using reverse transcriptase SuperScript III (Invitrogen) in the manufacturer's reaction buffer for 1 hour at 55°C. Remaining RNA template was degraded by addition of NaOH to a final concentration of 50 mM and heating at 80°C for 10 minutes, followed by quenching with HCl. The final samples were diluted in formamide loading buffer and separated on 8% dPAGE gels containing 7 M urea. Bands were visualized using the LI-COR Odyssey Imaging System.

**Western Blotting.** Cells were harvested by scraping into ice-cold PBS solution, and lysed in RIPA buffer (50 mM Tris pH 7.4, 150 mM NaCl, 1 mM EDTA, 1% Nonidet P-40, 0.1% Na-deoxycholate, 0.1% SDS, protease inhibitor tablet (Roche), 1 mM PMSF). Proteins were separated in 8-12% SDS-PAGE gels and transferred to nitrocellulose membranes. Membranes were then blocked with 5% BSA in TBST (5 mM Tris-HCl, pH 7.5, 15 mM NaCl, 1% Tween-20) before application of the indicated antibodies (Table S1). After incubation with IR-labeled secondary antibodies and washes in 1×TSBT, blots were imaged on a LI-COR Odyssey Imaging System.

**Cell Viability Assays.** Cells were plated in 96-well culture plates (5,000 cells in 200 µL of medium per well) 16 hours prior to treatment with various concentrations of the indicated drug compounds. Cell viability after 72 hours was measured using the CellTiter 96 Aqueous One Solution Cell Proliferation Assay (MTS) (Promega) according to the manufacturer's instructions. Absorbance was measured at 490 nm using a Synergy H1 Microplate Reader (BioTek), and cell viability was calculated as the ratio of absorbance of treated cells to that of untreated cells.

**Statistical Analysis.** Graphs were generated with Prism 9 for Windows (GraphPad Software Inc., Version 9.3.1). Unless otherwise indicated, horizontal bars represent the mean values and error bars the standard deviation. In box plot graphs, boxes represent the 25-75 percentile range, and whiskers represent the 9-95 percentile range. Data points outside this range are shown individually. Statistical tests were applied as described in the Figure legends and were calculated using Prism 9.

**Table S1.** Antibodies used in this study.

| Antibody | Clone No. | Company | Catalogue No. | Application / Dilution |
|---|---|---|---|---|
| NPM1 | FC-61991 | Thermo Fisher (Invitrogen) | 32-5200 | IF 1:300 |
| TCOF1 | --- | Millipore Sigma | HPA038237 | IF 1:150 |
| p53 | DO-1 | Millipore Sigma | OP43 | WB 1:1000 |
| β-Actin | 8H10D10 | Cell Signaling | 3700 | WB 1:10,000 |
| UBF | F-9 | Santa Cruz | sc-13125 | WB 1:1000 ; IF 1:250 |
| RPA194 | C-1 | Santa Cruz | sc-48385 | WB 1:500 ; IF 1:100 |
| TAF I p110 | C-10 | Santa Cruz | sc-374551 | WB 1:500 |
| phospho-H2AX (S193) | JBW301 | Millipore Sigma | 05-636 | IF 1:400 |
| NBS1 | --- | Novus Biologicals | NB100-143 | IF 1:150 |
| TOPBP1 | --- | Thermo Fisher (Bethyl Laboratories) | A300-111A-M | IF 1:200 |
| MRE11 | 12D7 | GeneTex | GTX70212 | IF 1:100 |
| phospho-RPA (S4,S8) | --- | Thermo Fisher (Bethyl Laboratories) | A300-245A | IF 1:100 |
| FLAG | M2 | Millipore Sigma | F1804 | IF 1:1000 |
| HA | 6E2 | Cell Signaling | 2367 | IF 1:1000 |
| CHK1 | G-4 | Santa Cruz | sc-8408 | WB 1:1000 |
| phospho-CHK1 (S345) | 133D3 | Cell Signaling | 2348 | WB 1:500 |
| CHK2 | --- | Cell Signaling | 2662 | WB 1:1000 |
| phospho-CHK2 (T68) | C13C1 | Cell Signaling | 2197 | WB 1:2000 |

**Table S2.** siRNA molecules used in this study.

| Name | Sequence (5' to 3') |
|---|---|
| siCtrl | UGGUUUACAUGUCGACUAA |
| siTOPBP1 | ACAAAUACAUGGCUGGUUA |
| siTCOF1 | CCACCAUGGGUUGGAACUAAAUU |

**Table S3.** Primers for rDNA amplification and rRNA primer extension.

| Name | Sequence (5' to 3') |
|---|---|
| prHu3-Promoter Forward | Biotin-CTCCCGCGTGTGTCCTGGGGTTGACCAGAG |
| prHu3-Promoter Reverse | GGCACGGTGGCCCTCGCCGCCTTCC |
| 28S-rRNA-RT (nt 4455-4479) | IRDye800-GCTCTTCCTATCATTGTGAAGCAGA |
| 18S-rRNA-RT (nt 961-992) | IRDye800-TCTGGTCCGTCTTGCGCCGGTCCAAGAATTT |
| 5S-rRNA-RT (nt 101-121) | IRDye800-AAAGCCTACAGCACCCGGTAT |

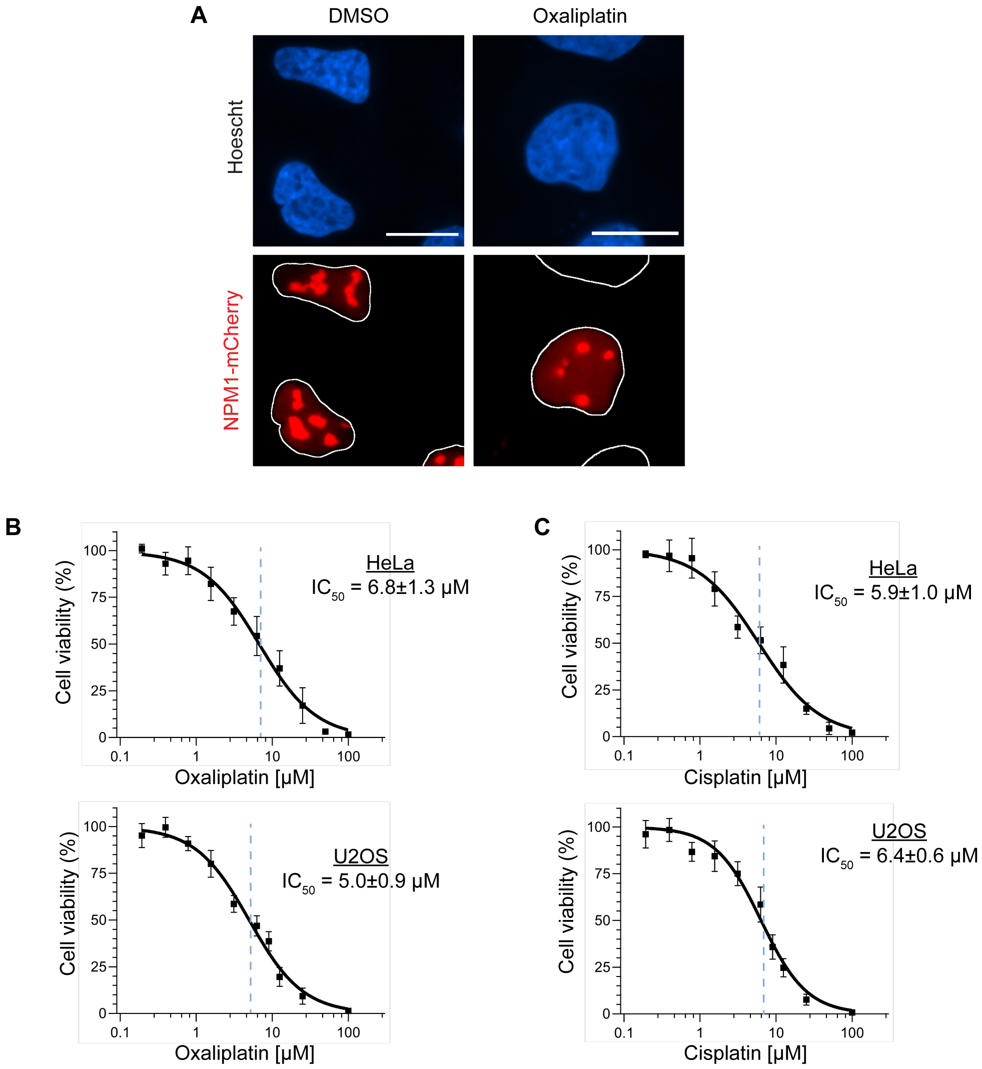


**Figure S1.** *Sensitivity of cancer cell lines to Pt drugs and localization of NPM1-mCherry.* (A) Representative images of NPM1-mCherry localization in wild type U2OS cells after 6 hours of treatment with 5 μM oxaliplatin. White outlines of nuclei were generated from the DAPI channel. (B-C) Dose response curves for oxaliplatin (B) and cisplatin (C) in HeLa and U2OS cell lines. Cell viability was measured using MTS assay at 72 hours post-treatment and corrected for DMSO treatment control. Dose response curves were generated via nonlinear fit to the data, and IC50 values were derived from the fits. Error bars represent mean ± SEM of *N* = 3 biological replicates. All scalebars = 15 μM.

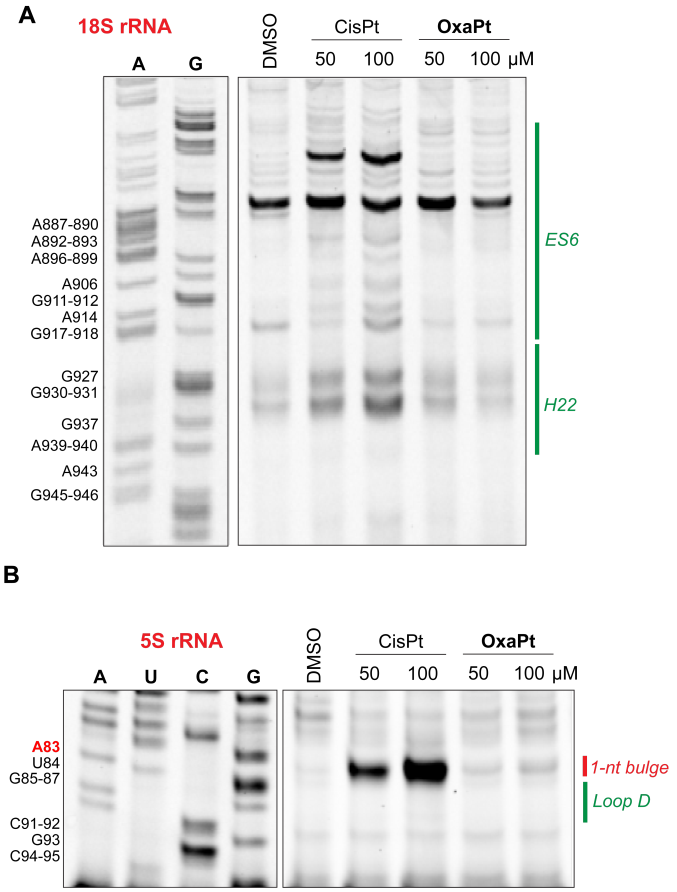


**Figure S2.** *Comparison of platinum adduct formation on cellular rRNA.* Primer extension assays of 18S (A) and 5S (B) human rRNA extracted from U2OS cells treated with 0-100 µM of cisplatin or oxaliplatin for 12 hours. Dideoxy sequencing ladder labeling A, U, C, and/or G is shown on the left. Annotations of local rRNA structures are provided on the right.

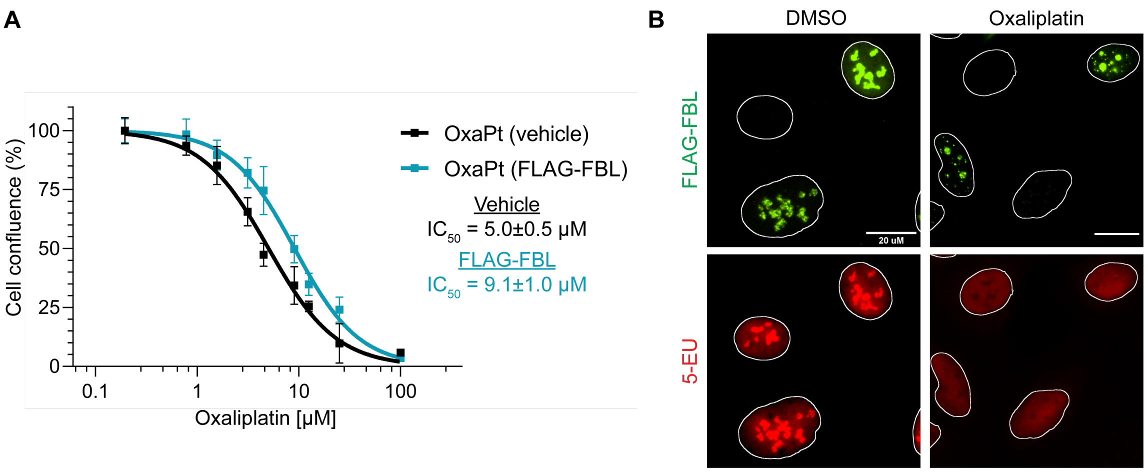


**Figure S3.** *Overexpression of FBL does not rescue transcriptional silencing by oxaliplatin.* (A) Dose response curves for oxaliplatin in U2OS cells after transient transfection with FLAG-FBL. Medium containing oxaliplatin was added 24 hours after transfection of the DNA plasmid, and cell viability was measured using MTS assay at 72 hours after addition of Pt drug and corrected for DMSO treatment control in each transfection condition. Dose response curves were generated via nonlinear fit to the data, and IC50 values were derived from the fits. Error bars represent mean ± SEM of $N$ = 3 biological replicates. (B) Localization of transiently transfected FLAG-FBL in U2OS cells and measurement of rRNA synthesis by 5-EU incorporation after treatment with 5 μM oxaliplatin for 6 hours. White outlines of nuclei were generated from the DAPI channel. All scalebars = 20 μM.

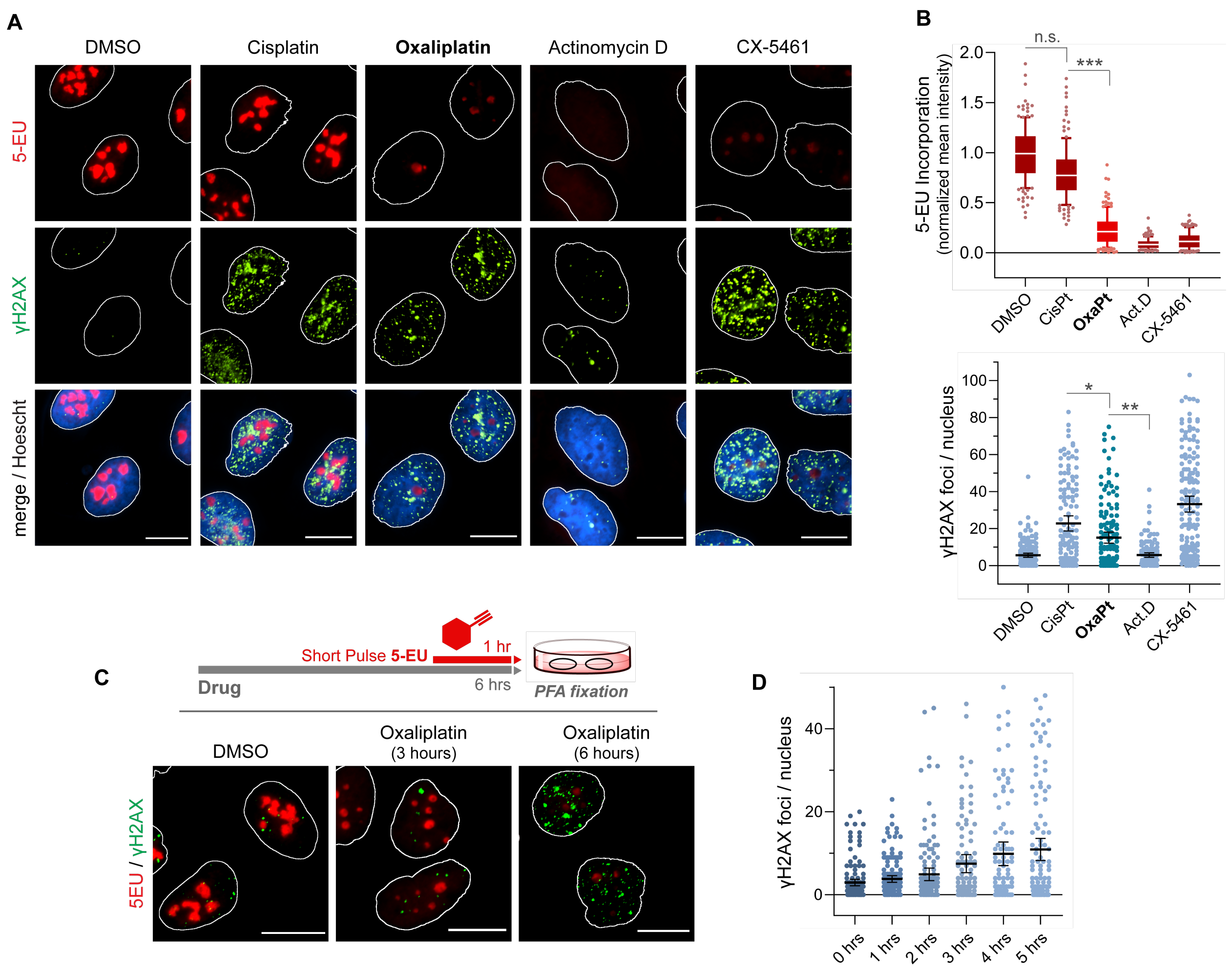


**Figure S4.** *Oxaliplatin induces a modest γH2AX response.* (A) Representative images of immunostaining against phospho-H2AX (γH2AX) and measurement of nucleolar RNA synthesis by 5-EU incorporation after treatment of U2OS cells with cisplatin (5 μM), oxaliplatin (5 μM), Actinomycin D (5 nM), or CX-5461 (1 μM) for 6 hours. (B) Quantification of changes in nucleolar 5-EU incorporation in response to drug treatments, relative to DMSO control *(top)*, and quantification of γH2AX foci count per nucleus, measured in ImageJ using ROIs from DAPI channel *(bottom)*. Number of cells analyzed for each condition, summed over 3 biological replicates: DMSO (*N* = 151), cisplatin (*N* = 135), oxaliplatin (*N* = 142), Actinomycin D (*N* = 121), CX-5461 (*N* = 164). Error bars represent mean ± SEM. Statistical analysis in (B) was performed using a one-way analysis of variance, Tukey's multiple comparisons test. Adjusted *p*-values are indicated by ***P<0.001, **P<0.01, *P<0.05, and *n.s.* denotes a non-significant *p*-value > 0.05. (C) Schematic of "short-pulse" 5-EU labeling of nucleolar RNA synthesis, and representative images of fixed cells at key time-points for γH2AX induction and rRNA silencing upon treatment with oxaliplatin (5 μM). (D) Quantification of γH2AX foci from time-course in (C). Number of cells analyzed at each time point post-treatment, summed over 3 biological replicates: 0 hours (*N* = 96), 1 hour (*N* = 104), 2 hours (*N* = 126), 3 hours (*N* = 155), 4 hours (*N* = 98), 5 hours (*N* = 133). In (A) and (C), white outlines of nuclei were generated from the DAPI channel. All scalebars = 15 μM.

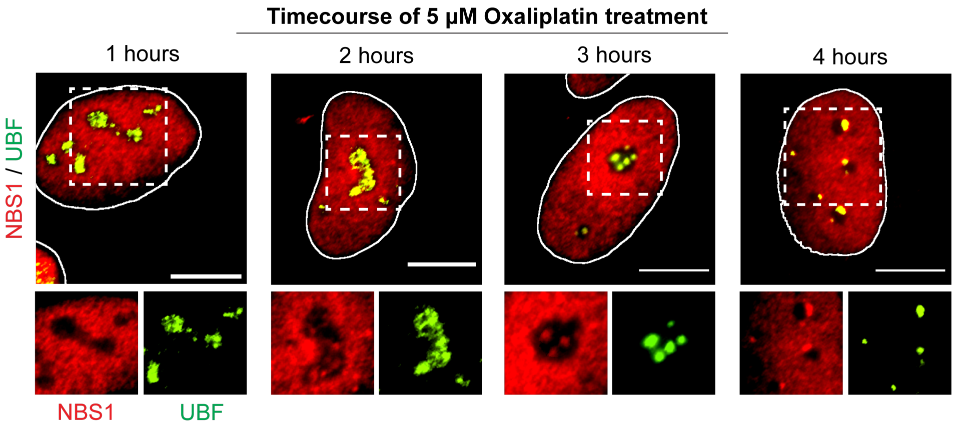


**Figure S5.** *Time-course of NBS1 recruitment to nucleolar caps.* (A) Representative images of fixed U2OS cells at specified time-points immunostained against NBS1 (red) and UBF (green) after treatment with 5 µM oxaliplatin. Hashed box indicates region of zoom for cropped single-channel images below. White outlines of nuclei were generated from the DAPI channel, and all scalebars = 15 µM.

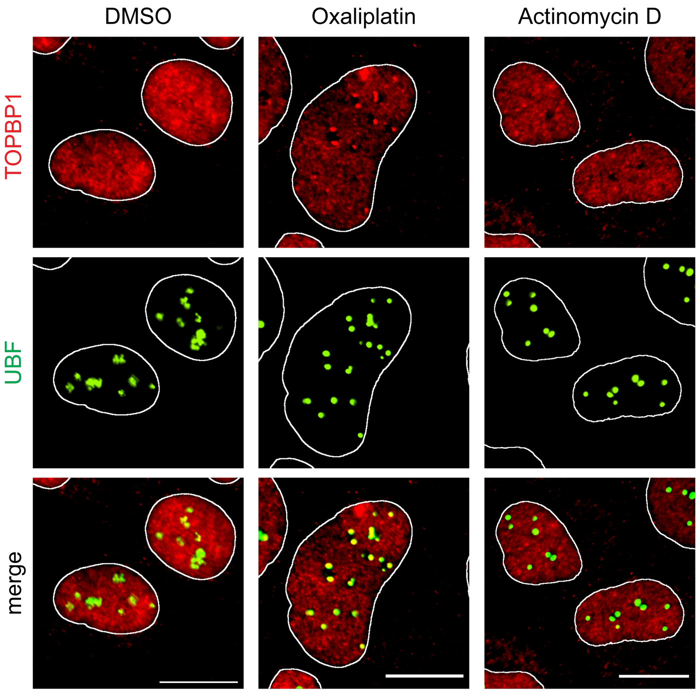


**Figure S6.** *TOPBP1 is recruited to nucleoli upon oxaliplatin treatment.* Representative images of immunostaining against TOPBP1 (red) and UBF (green) after treatment with oxaliplatin (5 μM) and Actinomycin D (5 nM) for 6 hours. White outlines of nuclei were generated from the DAPI channel, and all scalebars = 15 μM.

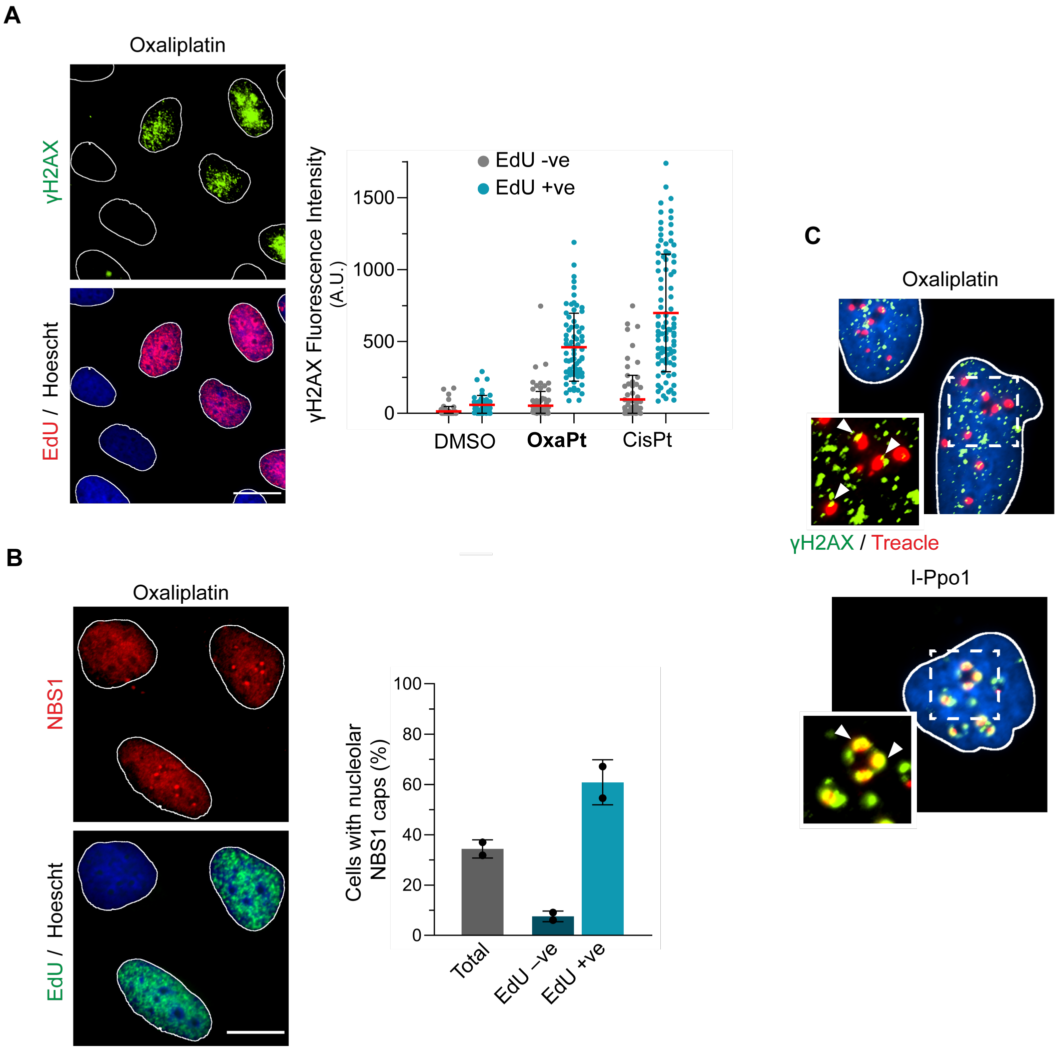


**Figure S7.** *NBS1 is recruited to nucleoli of S-phase cells exhibiting DNA damage signaling.* (A) Representative images of immunostaining against γH2AX and labeling of EdU incorporation into cells treated with 5 μM oxaliplatin for 6 hours *(left)*, and quantification of γH2AX signal intensity in EdU-positive and -negative cells after treatment with oxaliplatin or cisplatin *(right)*. (B) Representative images of immunostaining against NBS1 (red) and labeling of EdU (green) in cells treated with 5 μM of oxaliplatin *(left)*. Fraction of EdU-positive and EdU-negative cells with nucleolar caps containing NBS1, sum of 2 biological experiments *(right)*. Error bars in all graphs represent mean ± SEM. All experiments were performed in wild-type U2OS cells. All scalebars = 15 μM. (C) Representative images of immunostaining against γH2AX (green) and Treacle (red) in cells treated with oxaliplatin (5 μM) or transiently transfected with I-Ppo1 for 8 hours. Merged images with Hoescht nuclear staining are shown, with hashed boxes indicating region of zoom for cropped images highlighting γH2AX/Treacle colocalization around nucleoli.

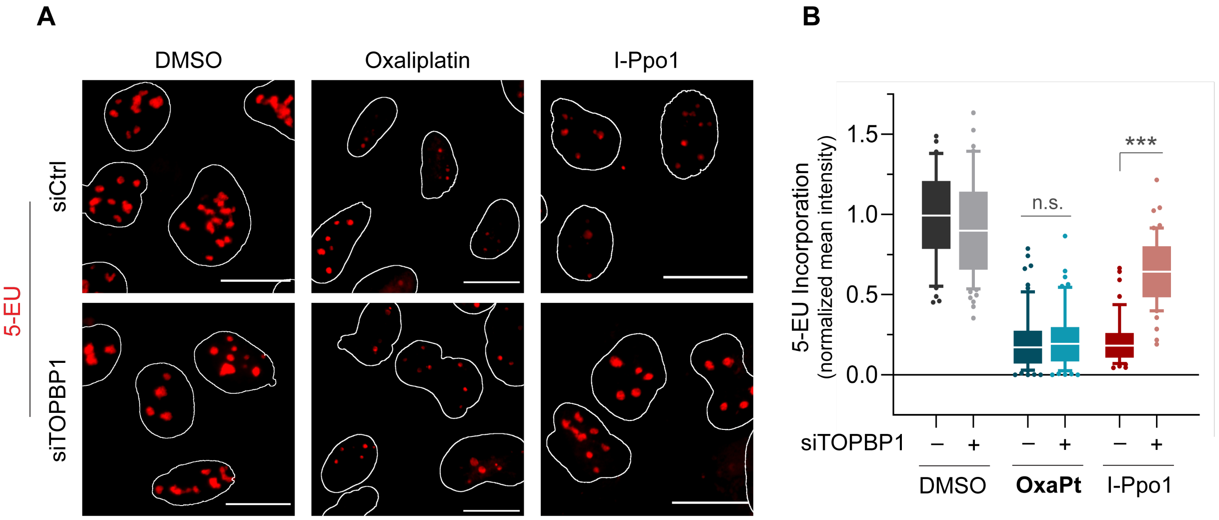


**Figure S8.** *TOPBP1 is not essential for rRNA silencing by oxaliplatin.* (A) Measurement of rRNA synthesis by 5-EU incorporation after treatment with oxaliplatin (5 μM, 6 hours) or transfection of I-Ppo1 (8 hours) in siCtrl and siTOPBP1-treated cells. White outlines of nuclei in were generated from the DAPI channel, and all scalebars = 15 μM. (B) Quantification of 5-EU incorporation from (A), relative to DMSO control. Number of cells analyzed in each condition, summed over 2 biological replicates: DMSO in siCtrl ($N$ = 43) and siTOPBP1 ($N$ = 72), OxaPt in siCtrl ($N$ = 64), and siTOPBP1 ($N$ = 59), I-Ppo1 in siCtrl ($N$ = 46) and siTOPBP1 ($N$ = 50). For I-Ppo1-treated cells, only cells positive for HA staining were quantified. Statistical analysis was performed using a one-way analysis of variance, Tukey's multiple comparisons test. Adjusted p-value is indicated by ***P<0.001, and *n.s.* denotes a non-significant $p$-value > 0.05.
